# The enteric nervous system of *C. elegans* is specified by the Sine Oculis-like homeobox gene *ceh-34*

**DOI:** 10.1101/2021.11.30.470650

**Authors:** Berta Vidal, Burcu Gulez, Wen Xi Cao, Eduardo Leyva-Diaz, Tessa Tekieli, Oliver Hobert

## Abstract

Overarching themes in the terminal differentiation of the enteric nervous system, an autonomously acting unit of animal nervous systems, have so far eluded discovery. We describe here the overall regulatory logic of enteric nervous system differentiation of the nematode *C. elegans* that resides within the foregut (pharynx) of the worm. A *Caenorhabditis elegans* homolog of the *Drosophila* Sine Oculis homeobox gene, *ceh-34*, is expressed in all 14 classes of interconnected pharyngeal neurons from their birth throughout their life time, but in no other neuron type of the entire animal. Constitutive and temporally controlled *ceh-34* removal shows that *ceh-34* is required to initiate and maintain the neuron type-specific terminal differentiation program of all pharyngeal neuron classes, including their circuit assembly, without affecting panneuronal features. Through additional genetic loss of function analysis, we show that within each pharyngeal neuron class, *ceh-34* cooperates with different homeodomain transcription factors to individuate distinct pharyngeal neuron classes. Our analysis underscores the critical role of homeobox genes in neuronal identity specification and links them to the control of neuronal circuit assembly of the enteric nervous system. Together with the pharyngeal nervous system simplicity as well as its specification by a Sine Oculis homolog, our findings invite speculations about the early evolution of nervous systems.

## INTRODUCTION

Across animal phylogeny, the enteric nervous system constitutes a self-contained, autonomously acting neuronal network that detects physiological conditions to control the peristaltic movement of food through the digestive tract (Ayali, 2004; Copenhaver, 2007; Fung and Vanden Berghe, 2020; Hartenstein, 1997; Laranjeira and Pachnis, 2009). Because of structural and functional autonomy, the enteric nervous system of humans has been referred to as a “second brain” (Gershon, 1998). In mammals, the enteric nervous system is composed of around 20 different neuron types, categorized into internal sensory, inter- or motor neurons, which line the interior lumen of different sections of the digestive system (Drokhlyansky et al., 2020; Furness, 2000). While a lot of progress has been made in understanding early developmental patterning events that establish the fate of neurons in the enteric nervous system of mammals (Nagy and Goldstein, 2017; Sasselli et al., 2012), fish (Ganz, 2018) and flies (Copenhaver, 2007; Myers et al., 2018), very little is known about the later steps of enteric neurons terminal differentiation, both in vertebrate and invertebrate models (Copenhaver, 2007; Hao and Young, 2009; Memic et al., 2018; Morarach et al., 2021; Rao and Gershon, 2018). Specifically, it has remained unclear as to whether there are common unifying themes in how enteric neurons acquire their terminally differentiated state. This is particularly interesting from an evolutionary standpoint. The function of enteric neurons in controlling feeding behavior is an ancient one that may precede the evolution of the bilaterian central nervous system (Cook et al., 2020; Furness and Stebbing, 2018; Gilbert, 2019; Koizumi, 2007). Understanding how enteric neurons acquire their terminal features may therefore provide novel insights into nervous system evolution.

The nematode *C. elegans* contains an enteric nervous system in its foregut, the pharynx (Albertson and Thomson, 1976; Cook et al., 2020; Mango, 2007). The pharyngeal nervous system shares features of enteric nervous systems of more complex animals. It is an autonomously acting unit, composed of 20 synaptically interconnected neurons that fall into 14 anatomically distinct classes, required for movement of food through the digestive tract of the worm (**Fig.1**)(Albertson and Thomson, 1976; Avery, 2012; Cook et al., 2020; Mango, 2007). Like other enteric nervous systems, the nematode pharyngeal nervous system is a non-centralized neuronal network isolated from the rest of the nervous system and, in rough analogy to the vagus nerve, is connected to the remainder of the nervous system through a single nerve fiber, that of the bilateral RIP neuron pair (Albertson and Thomson, 1976; Cook et al., 2020; Cook et al., 2019; White et al., 1986). Like vertebrate enteric neurons (Fung and Vanden Berghe, 2020), pharyngeal neurons have sensory, inter- and motorneuron function (Avery and Horvitz, 1989; Cook et al., 2020; Trojanowski et al., 2014). Our recent re-analysis of pharyngeal nervous system anatomy has shown that rather than segregating these functions over distinct neurons as vertebrate do (Fung and Vanden Berghe, 2020), most pharyngeal neurons each combine sensory, inter- and motorneuron function, i.e. are polymodal (Cook et al., 2020). The 14 distinct types of pharyngeal neurons are defined by their unique anatomy, i.e. axonal projections, morphology, synaptic connectivity (**Fig.1B,C**)(Cook et al., 2020) and unique functional features (Avery and Horvitz, 1989; Trojanowski et al., 2014). This anatomical and functional classification has recently been further extended by the description of their unique molecular fingerprint, determined by scRNA transcriptomic analysis (Taylor et al., 2021)(**Fig.1D**). For example, each pharyngeal neuron class is uniquely defined by characteristic signatures of neurotransmitter systems, from acetylcholine (ACh), glutamate (Glu) to serotonin, and by unique combinations of neuropeptide-encoding genes (**Fig.1E**). While pharyngeal neurons are clearly different from one another, scRNA profiling has shown that their molecular signatures are more similar to each other than to other neurons in the nervous system (Taylor et al., 2021)(**Fig.1D**).

**Figure 1:**
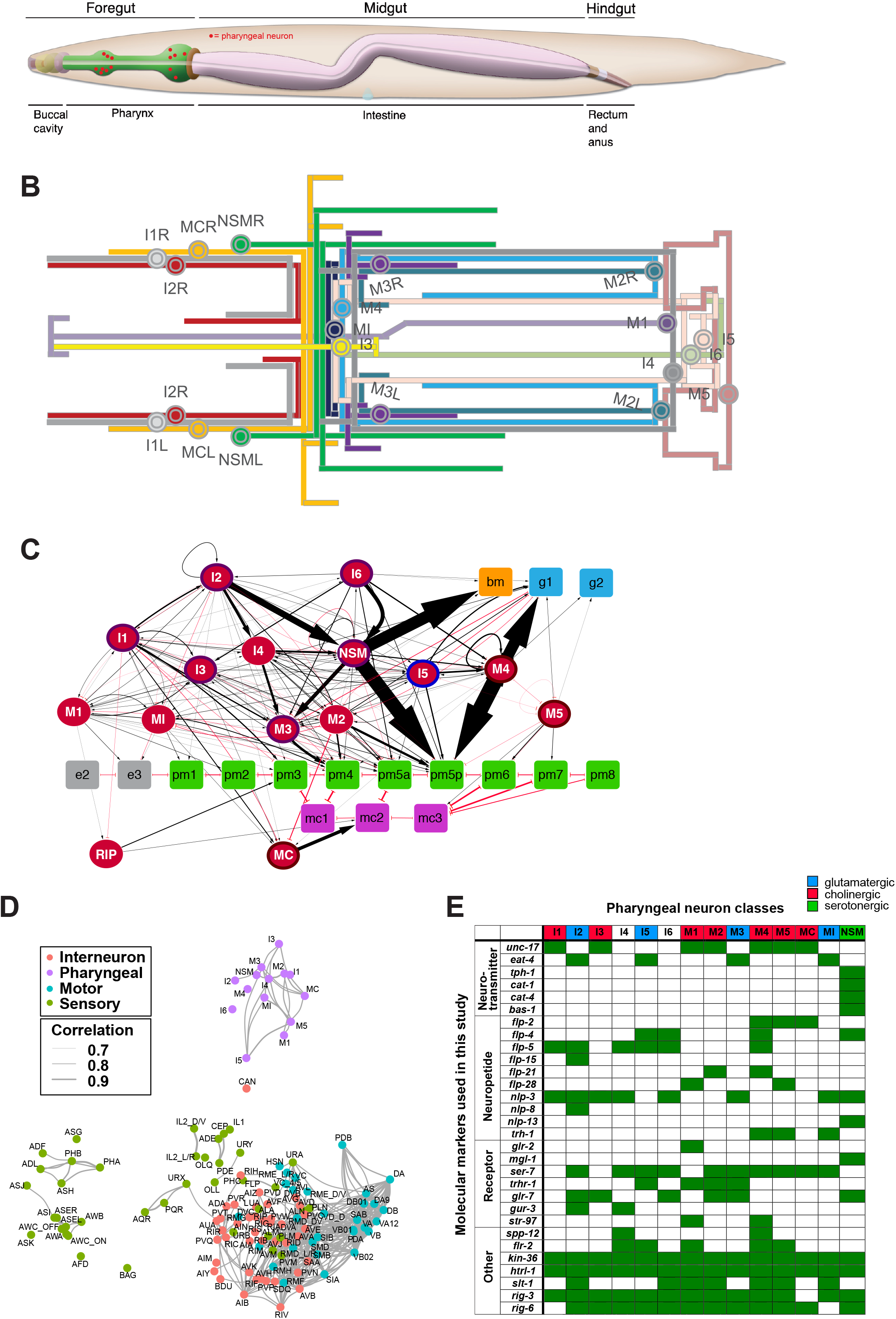
The pharyngeal nervous system of *C. elegans*. **(A)** Overview of alimentary system of *C. elegans* from Wormbook (Hall and Altun, 2007), with neuronal cell bodies in the pharynx added in red. **(B)** Projection patterns of pharyngeal neurons within the pharynx displayed in the format of a subway map (kindly provided by S.J. Cook). . **(C)** Full connectome of pharyngeal nervous system, adapted from (Cook et al., 2020). Square nodes are end-organs, including muscle (green), marginal cells (fuchsia), gland cells (blue), epithelial cells (grey), and basement membrane (orange). Neurons are red ellipses. Neurons with outlines have either apical (purple), unexposed (brown), or embedded (blue) sensory endings. Directed chemical edges and undirected gap junction edges are represented by black arrows and red lines, respectively. The line width is proportional to the anatomical strength of that connection (# serial sections). The pharyngeal nervous system is connected to the rest of the nervous system through a single neuron pair (RIP). **(D)** Single cell transcriptome similarity between neuron types classes with widths of edges indicating strengths of similarity (Pearson correlation coefficients >0.7), showing that pharyngeal neurons are more similar to each other than to other neurons in the *C. elegans* nervous system. Reproduced from (Taylor et al., 2021). **(E)** Molecular markers used in this study for cell fate analysis. See **Supplemental File 1** for information on reporter constructs.

One fascinating aspect of the *C. elegans* pharyngeal nervous system is that it has an appearance of what one could imagine an ancestral, primitive nervous system to have looked like. Primitive nervous systems are generally thought to have emerged out of the context of monolayers of epithelial cells, with individual cells in such layers specializing into primitive sensory-motor-type neurons (Arendt, 2008; Mackie, 1970). These diffusely organized neurons may have sensed the environment and relayed such sensory information to the other primitive cell type thought to have arisen early in evolution, namely, contractile “myoepithelial” cells that were able to generate motion. The organization of the *C. elegans* pharynx reveals some striking parallels to such a presumptive primitive nervous system: It is also essentially a single monolayer of cells that are organized into a tubular structure and the vast majority of constituent cells are myoepithial cells (pharyngeal muscle) and polymodal, interconnected sensory/motor neurons (Cook et al., 2020; Mango, 2007; Portereiko and Mango, 2001). Pharyngeal neurons combine sensory, inter- and motor neuron features and are also not localized to ganglia but rather diffusely localized, resembling the architecture of more ancient nerve nets (Albertson and Thomson, 1976; Watanabe et al., 2009). The interconnectivity of pharyngeal neurons displays also less selectivity that non-pharyngeal neurons, such that mere physical proximity is an almost sufficient criteria for connectivity (Cook et al., 2020). Moreover, pharyngeal neurons are closely related to non-neuronal pharyngeal cells by lineage; for example, some muscle and neurons derive from a single mother cells (Sulston et al., 1983). The idea of enteric neurons being reflective of an early, primitive state of the nervous system has also been brought forward in the context of comparing enteric nervous systems from widely divergent species (Furness and Stebbing, 2018; Gilbert, 2019). Specifically, these authors argued that the nerve net-like hydra nervous system displays features of the vertebrate enteric nervous system.

The self-contained and hypothetically primitive state of the *C. elegans* enteric nervous system, with all its unique features, encouraged us to use this system as a model to probe several concepts of neuronal identity specification that have emerged from the centralized, non-pharyngeal nervous system of *C. elegans*: (1) The first is the concept of terminal selectors, transcription factors that act in a master-regulatory manner to coordinately control the many identity features of a terminally differentiating neuron (Hobert, 2016). Are members of terminal gene batteries in each pharyngeal neuron also controlled in a coordinated manner, via terminal selectors? One study in the NSM neurons provided some limited evidence in this regard (Zhang et al., 2014), but how broadly this applies throughout the pharyngeal nervous system was less clear. (2) Second, homeodomain transcription factors have a predominant role as terminal selectors of neuronal identity in the non-pharyngeal nervous system (Hobert, 2021; Reilly et al., 2020). A recent cataloguing of the expression patterns of all homeodomain proteins in the *C. elegans* genome revealed that each one of the *C. elegans* 118 neuron classes, including the pharyngeal neurons, display a unique combinatorial signature of homeobox gene expression (Reilly et al., 2020). Are all pharyngeal neurons indeed specified by homeobox gene combinations(s)? (3) Lastly, there is evidence from both *C. elegans* (Berghoff et al., 2021; Pereira et al., 2015) and other systems (Brunet and Pattyn, 2002), that synaptically connected neurons are often specified by the same transcription factor, suggesting that such transcription factors may be involved in assembling neurons in functional circuitry. We have termed such transcription factors “circuit organizers” (Berghoff et al., 2021; Pereira et al., 2015) and sought to test whether the isolated pharyngeal circuitry is similarly specified by a circuit organizer transcription factor.

In this paper, we show that all three predictions are fulfilled in the context of the nematode’s pharyngeal/enteric nervous system. We show that a *C. elegans* ortholog of the Sine Oculis/Six1/Six2 homeobox gene, *ceh-34*, is expressed in all pharyngeal neurons from their birth throughout their life time. Its expression is induced by the foregut organ selector gene *pha-4/FoxA*. We demonstrate that *ceh-34* initiates and maintains the terminally differentiated state of all synaptically connected pharyngeal neurons, that *ceh-34* is required to assemble pharyngeal neurons into proper circuitry and that in distinct pharyngeal neuron types, *ceh-34* cooperates with distinct homeobox genes to specify their respective identity. Taken together, our studies further substantiate overarching themes of nervous system development and potentially provide insights into the evolution of nervous systems.

## RESULTS

### Expression of paralogous genes *ceh-34* and *ceh-33*, the two *C. elegans* Sine Oculis/Six1/2 orthologs

The first systematic analysis of developmental patterning genes conducted after the release of the *C. elegans* genome sequence revealed several *C. elegans* homologs of the Sine oculis/Six family of homeodomain proteins (Ruvkun and Hobert, 1998) whose founding member was first identified in *Drosophila* for its role in eye patterning (Cheyette et al., 1994). This specific homeodomain transcription factor family is characterized by the presence of a conserved domain, located N-terminally to the DNA binding homeodomain, the ∼150 amino acid long SIX domain, involved in both protein-DNA, as well as protein-protein interactions (Kumar, 2009; Patrick et al., 2013). *C. elegans* Six-type homeodomain proteins fall into several, phylogenetically conserved families, the Sine Oculis/Six1/2, the Six4/5 and the Six3/6 subfamily (Dozier et al., 2001; Kumar, 2009). We focus here on the Sine Oculis subfamily.

Through the analysis of multiple nematode genome sequences we found that nematodes generally contain a single ortholog of the Sine Oculis/Six1/2 subfamily of SIX homeodomain proteins, but in the *Caenorhabditis* genus this locus has duplicated into two immediately adjacent paralogs, *ceh-33* and *ceh-34* (**Fig.2A**, **Fig.2 – Supplement 1**). Using a fosmid-based reporter in which the *ceh-33* locus is tagged with *gfp*, we find that the CEH-33 protein shows no expression in the nervous system within or outside the pharynx at any developmental stage. The only observed expression was in a subset of head muscle cells (**Fig.2B**).

**Figure 2:**
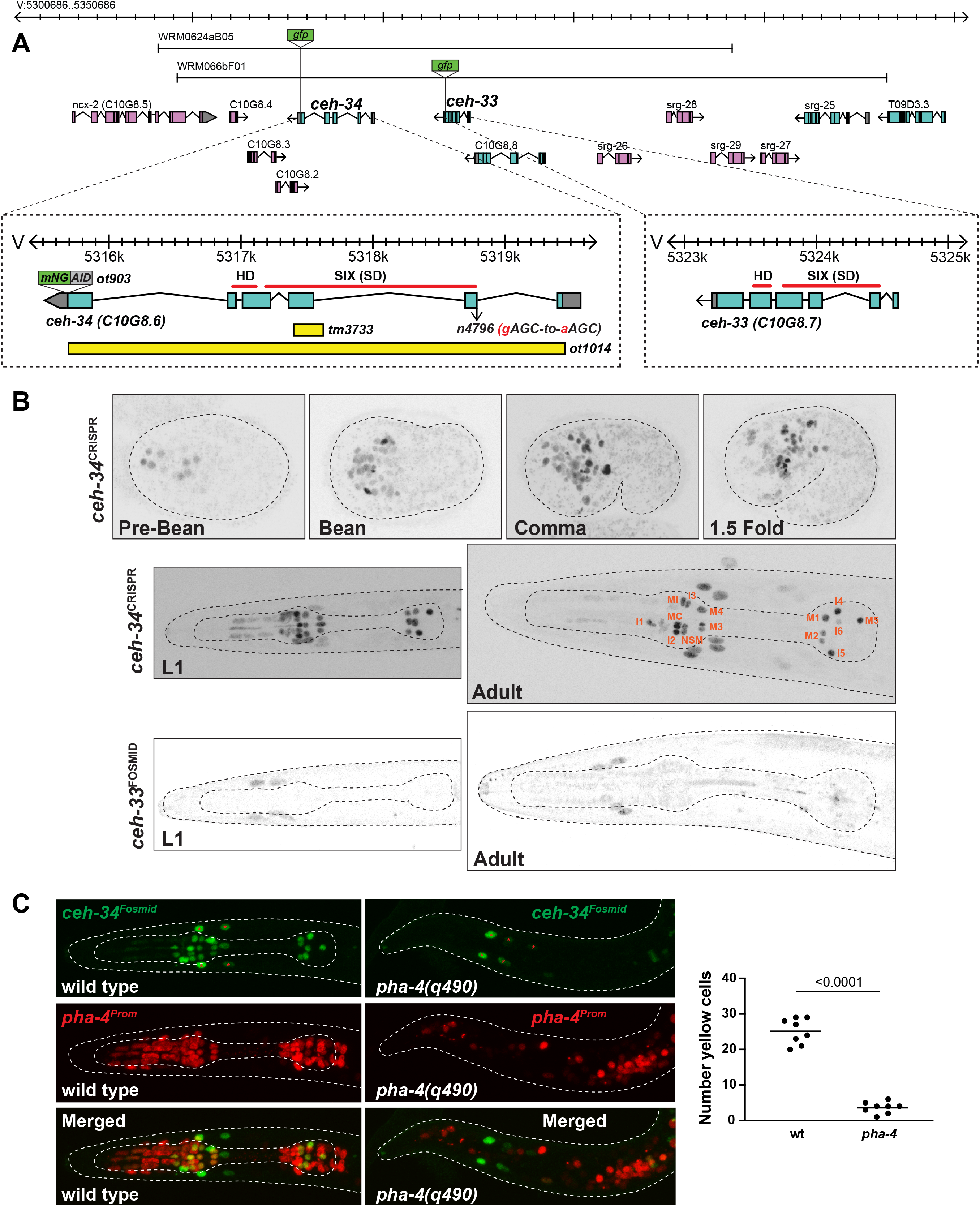
*ceh-34* is expressed in all pharyngeal neurons. **(A)** *ceh-33* and *ceh-34* loci showing different alleles and fosmid reporters used in this study. **(B)** Expression of the *ceh-34* CRISPR-engineered reporter allele *ot903* over the course of development. *ceh-33* fosmid reporter (*wgIs575*) shows expression in a subset of head muscle cells. **(C)** Pharynx organ selector *pha-4* controls *ceh-34* expression (as analyzed with the *wgIs524* transgene). Presumptive “pharyngeal cells” in *pha-4* mutant are marked with a red *pha-4* promoter fusion (*stIs10077*). Cells expressing *ceh-34* and *pha-4* were counted (yellow cells). *ceh-34* expression in head muscle cells, marked with red asterisk, is not affected since they do not express *pha-4*.

A previously described reporter construct that contains the coding region and 3.8kb of promoter region of the neighboring *ceh-34* gene showed expression in all pharyngeal neurons, but no other neurons outside the pharynx (Hirose et al., 2010). A fosmid-based reporter shows the same expression pattern (Reilly et al., 2020). To further confirm this strikingly selective pattern of neuronal expression and also to examine expression at different developmental stages with the best possible reagent, we used the CRISPR/Cas9 system to engineer a reporter allele of *ceh-34.* This reporter shows the same expression as the fosmid-based reporter in all pharyngeal neurons, but no other neurons (**Fig.2B**). The *ceh-34* reporter is turned on in the embryo at around the time of birth of pharyngeal neurons and is maintained in all pharyngeal neurons throughout all larval and adult stages (**Fig.2**).

The remarkable restriction of *ceh-34* expression within the nervous system to all pharyngeal neurons prompted us to ask whether the Forkhead transcription factor *pha-4*, an organ selector gene involved in early patterning of the pharynx (Gaudet and Mango, 2002; Horner et al., 1998; Kalb et al., 1998; Mango et al., 1994), is required for *ceh-34* expression. Crossing a *ceh-34* reporter into *pha-4(q490)* mutant animals, we indeed observed a loss of *ceh-34* expression (**Fig.2**). Conversely, *ceh-34* does not affect *pha-4* expression (**Fig.2** **– Supplement 2A)**. The regulation of *ceh-34* by *pha-4* mirrors the effects that *pha-4* has on the expression of other transcription factors that control terminal differentiation of other tissue types in the pharynx, for example the *ceh-22/NKX* homeobox gene that specifies pharyngeal muscle differentiation (Vilimas et al., 2004), or the *hlh-6* bHLH gene that specifies pharyngeal gland differentiation (Smit et al., 2008). We furthermore note that a *pha-4* reporter allele that we generated through CRISPR/Cas9 genome engineering is continuously expressed throughout the entire pharyngeal nervous system, at all postembryonic stages (**Fig.2** **– Supplement 2B**), raising the possibility that *pha-4* not may only initiate, but also maintain *ceh-34* expression.

### *ceh-34* controls the expression of diverse neurotransmitter signaling pathways in pharyngeal neurons

To begin to assess the function of *ceh-34* in enteric nervous system differentiation, we generated a null allele of the *ceh-34* locus, *ot1014,* using CRISPR/Cas9 engineering, in which the entire *ceh-34* locus is deleted (**Fig.2A**). Previously isolated alleles include a hypmorphic missense allele and a small deletion allele, *tm3733* (Amin et al., 2009; Hirose et al., 2010). In our ensuing mutant analysis, we found *ot1014* and the previously described strong loss of function allele *tm3733* are phenotypically indistinguishable and we therefore used both alleles interchangeably.

We first asked whether *ceh-34* is required for the generation of pharyngeal neurons. We found that in *ceh-34(tm3733)* mutant animals, the expression of the panneuronal genes *ric-4/SNAP25*, *ric-19/ICA1*, *rab-3/RAB3* and *unc-11/AP180* is unaffected (**Fig.3A**), indicating that pharyngeal neurons are generated and properly execute a generic differentiation program.

**Figure 3:**
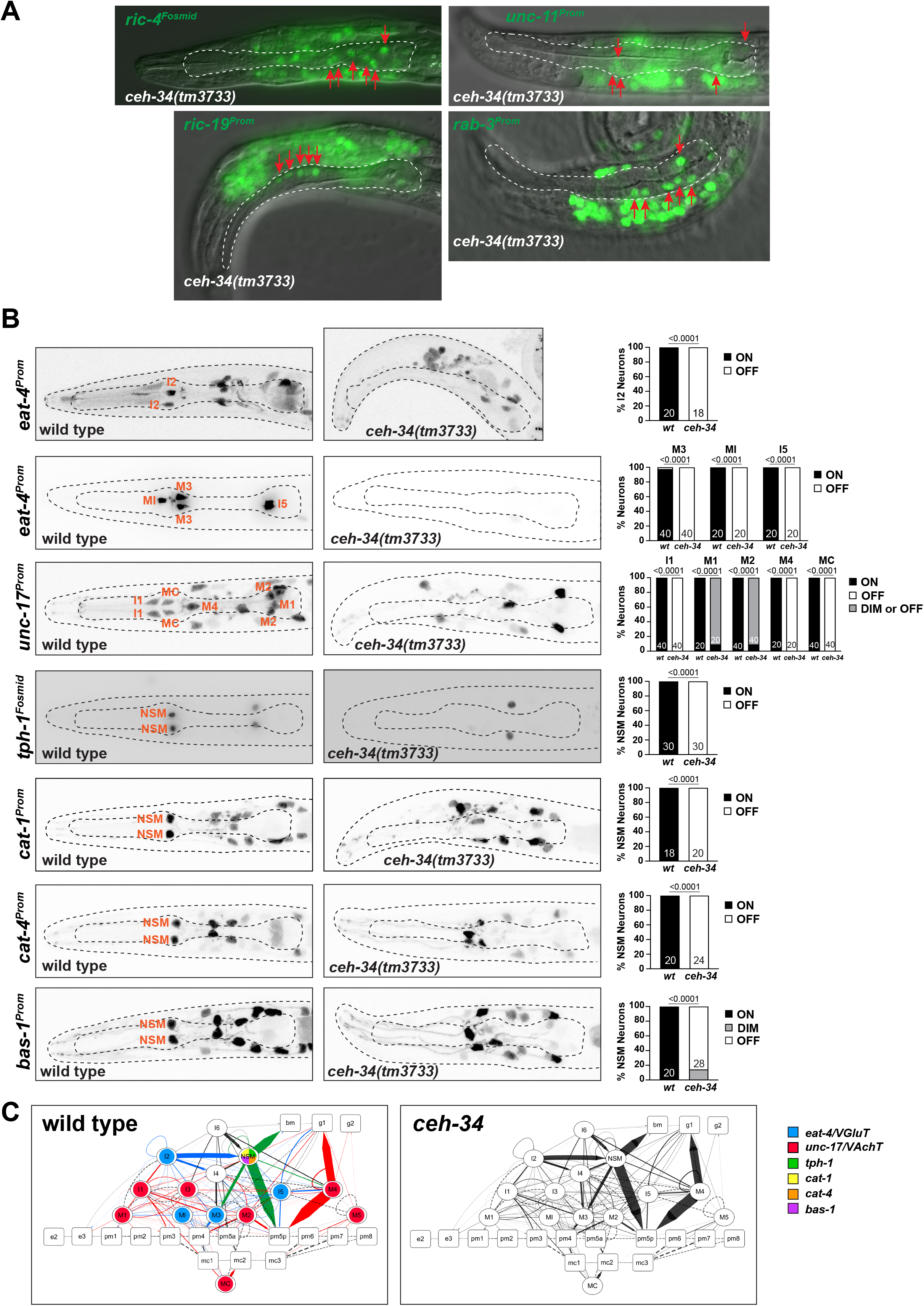
Pharyngeal neurons are generated in *ceh-34* mutants but loose their neurotransmitter identity. **(A)** Pictures showing expression of panneuronal reporter transgenes that monitor *ric-4 (otIs350)* , *ric-19 (otIs380)*, *unc-11 (otIs620)* and *rab-3 (otIs291)* expression. A single plane with a subset of pharyngeal neurons marked with red arrows is shown for clarity. **(B)** *ceh-34* affects the expression of neurotransmitter identity genes. Glutamatergic, cholinergic and serotonergic identity is lost. Reporter transgenes used are *eat-4 (otIs487, otIs558), unc-17 (otIs661), tph-1 (otIs517) , cat-1 (otIs221), cat-4 (otIs225) and bas-1 (otIs226).* Statistical analysis was performed using Fisher’s exact test or Chi-Square test. N is indicated within each bar and represents number of neurons scored. **(C)** Circuit diagram summarizing the effect of *ceh-34* on neurotransmitter identity. Nodes are colored to illustrate neurotransmitter identity gene expression. Nodes lose coloring when expression is affected in *ceh-34* mutants. Edges are colored if the source neuron expresses either *eat-4* (glutamatergic), *unc-17* (cholinergic) or *tph-1* (serotonergic). Edges lose coloring when expression of these genes is affected in the source neuron in *ceh-34* mutants (irrespective of whether the effect is partial or total). Note that in this and ensuing circuit diagrams, the existence of gray edges does not indicate whether those edges are generated properly in *ceh-34* mutants. Directed edges (arrows) represent chemical synapses. Undirected edges (dashed lines) represent electrical synapses. The width of the edge is proportional to the weight of the connection (the number of serial section electron micrographs where a synaptic specialization is observed).

We next assessed the effect of *ceh-34* on a large collection of neuron type-specific molecular identity features (illustrated in **Fig.1E**). To this end, we first focused on the signaling capacities of all neurons. In the vertebrate enteric nervous system, each individual neuron class is distinguished by its unique set of signaling molecules, from classic neurotransmitters to neuropeptides (Morarach et al., 2021). The same applies to all neuron classes in the pharyngeal/enteric nervous system of *C. elegans* (Horvitz et al., 1982; Pereira et al., 2015; Serrano-Saiz et al., 2013; Taylor et al., 2021). We first considered the three classic neurotransmitters ACh, Glu and serotonin which are employed in *C. elegans* much like in the enteric nervous system of other species: Seven of the 14 pharyngeal neuron classes use acetylcholine as neurotransmitter, four use glutamate and one uses serotonin (NSM) (Horvitz et al., 1982; Pereira et al., 2015; Serrano-Saiz et al., 2013; Taylor et al., 2021). Of those neurotransmitters, serotonin is perhaps the best studied neurotransmitter both in the vertebrate enteric nervous system (Gershon, 2013) as well as in the nematode pharyngeal nervous system (Horvitz et al., 1982; Ishita et al., 2020; Song and Avery, 2013). Acquisition of ACh, Glu and serotonin neurotransmitter identity features can be visualized through the expression of a number of enzymes and transporters: *unc-17/VAChT* for cholinergic identity, *eat-4/VGluT* for glutamatergic identity and *tph-1/TPH, cat-1/VMAT, bas-1/AAAD* and *cat-4/GCH* for serotonergic identity. We examined the expression of all these markers, using various reporter genes, in all pharyngeal neurons of *ceh-34* mutant animals and found that the seven cholinergic, four glutamatergic and single serotonergic neuron classes fail to acquire their respective neurotransmitter identity (**Fig.3B**). We schematize these results in the context of a pharyngeal circuit diagram to illustrate the breadth of effects of *ceh-34* on neurotransmitter identity acquisition (**Fig.3C**).

As in vertebrate enteric nervous systems (Drokhlyansky et al., 2020), pharyngeal neurons also display highly patterned expression of various neurotransmitter receptors (Taylor et al., 2021). We analyzed the *ceh-34-*dependence of three exemplary receptors, the serotonin receptor *ser-7*, which mediates several functions of pharyngeal serotonin (Hobson et al., 2006), as well as a metabotropic and an ionotropic Glu receptor, *mgl-1* and *glr-2,* each expressed in specific subsets of pharyngeal neurons. We find the expression of these receptors in different pharyngeal neuron types is strongly affected, if not entirely abrogated upon removal of *ceh-34* (**Fig.4**).

**Figure 4:**
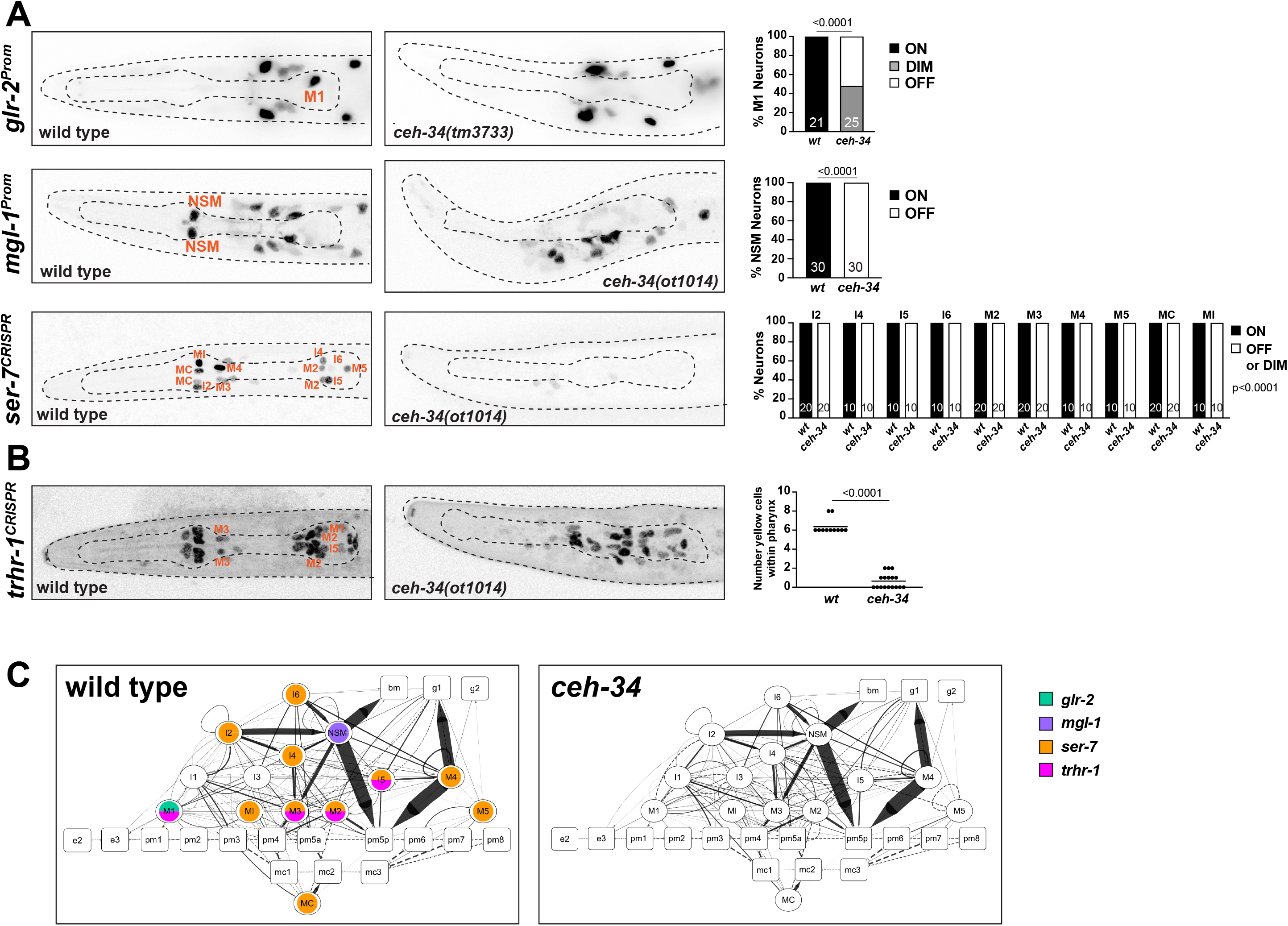
*ceh-34* affects the expression of receptors for neurotransmitters and neuropeptides. **(A)** Representative pictures and quantification showing neurotransmitter receptor expression loss in *ceh-34* mutants. Reporter genes used are reporter transgenes for *glr-2 (ivIs26), mgl-1 (otIs341)* and a CRISPR/Cas9-engineered reporter allele for *ser-7 (syb4502)*. Statistical analysis was performed using Fisher’s exact test or Chi-Square test. N is indicated within each bar and represents number of neurons scored. **(B)** Representative pictures and quantification showing neuropeptide receptor expression loss in *ceh-34* mutants. Reporter gene used is the CRISPR/Cas9-engineered reporter allele *trhr-1 (syb4453)*. Statistical analysis was performed using unpaired t-test. **(C)** Circuit diagram summarizing the effect of *ceh-34* on neurotransmitter and neuropeptide expression. Nodes lose coloring when expression is affected in a *ceh-34* mutant (irrespective of whether the effect is partial or total). See legend to Figure 3 for more information on features of circuit diagram.

### *ceh-34* controls diverse neuropeptidergic identities of pharyngeal neurons

The function of vertebrate enteric nervous system is modulated by a number of prominent neuropeptidergic signaling systems (Abot et al., 2018; Llewellyn-Smith, 1989). In some cases, both neuropeptide and receptor are expressed in the vertebrate enteric nervous system, while in others, the peptide is produced elsewhere but acts on neuropeptide receptors located in the enteric nervous system. Likewise, the *C. elegans* pharyngeal nervous system expresses a great diversity of neuropeptides and neuropeptide receptors, including homologs of neuropeptide signaling systems that function in the vertebrate enteric nervous system (Taylor et al., 2021). In fact, each pharyngeal neuron expresses a unique combination of neuropeptides and their receptor proteins (Taylor et al., 2021)(**Fig.5**). We tested whether *ceh-34* affects the neuron-type specific expression of various neuropeptidergic signaling systems.

**Figure 5:**
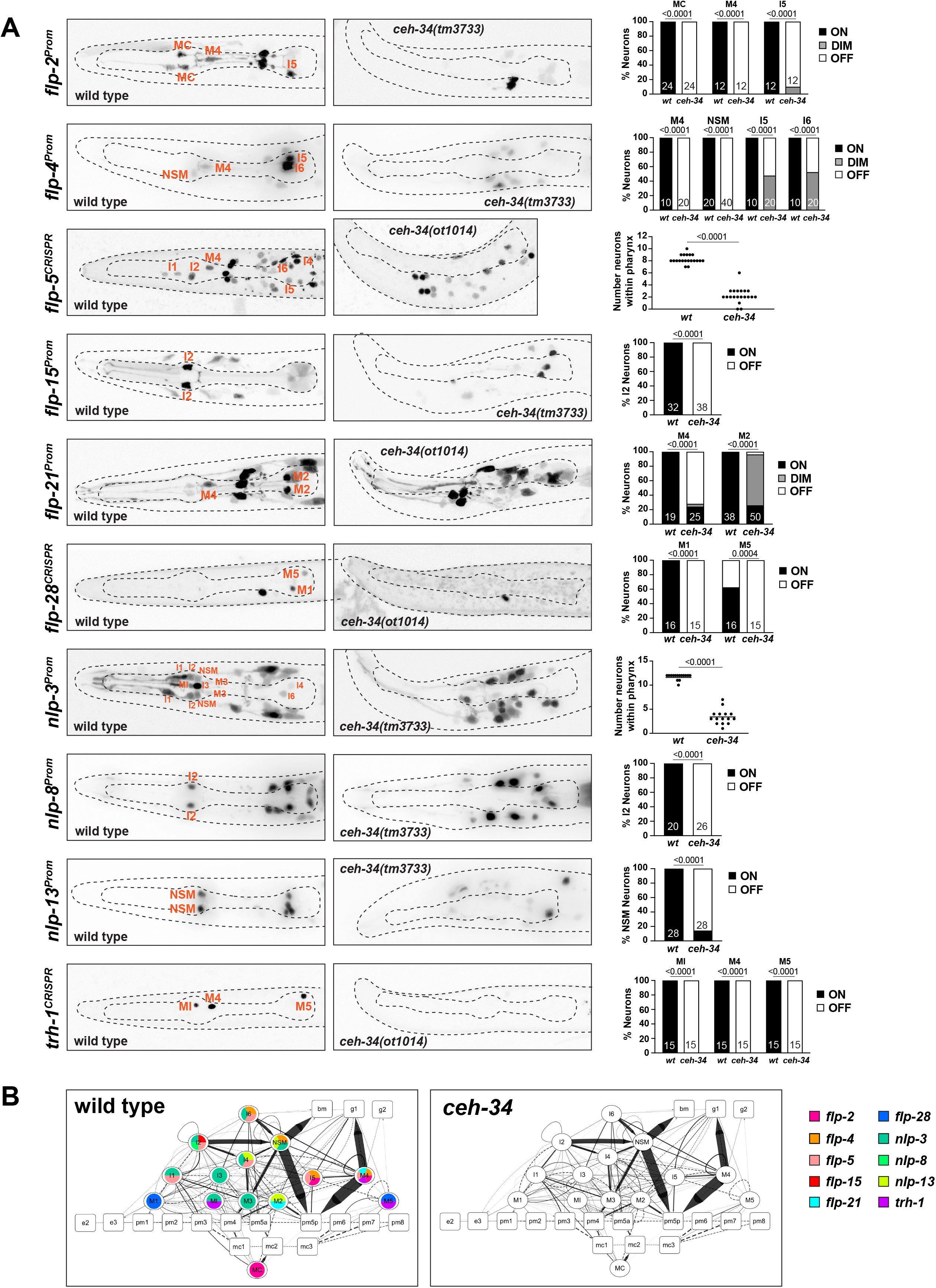
*ceh-34* affects neuropeptidergic identity of pharyngeal neurons. **(A)** Representative pictures and quantification showing expression of 10 different neuropeptides is affected in *ceh-34* mutants. Reporter genes used are transgenic reporters for *flp-2 (ynIs57), flp-4 (ynIs30), flp-15 (ynIs45), flp-21 (ynIs80), nlp-3 (otIs695), nlp-8 (otIs711)* and *nlp-13 (otIs742)* and CRISPR/Cas9-engineered reporter alleles for *flp-5 (syb4513), flp-28 (syb3207)* and *trh-1 (syb4421)*. Statistical analysis was performed using Fisher’s exact test, Chi-Square test or unpaired t-test. N is indicated within each bar and represents number of neurons scored. **(B)** Circuit diagram summarizing the effect of *ceh-34* on neuropeptide expression. Nodes lose coloring when neuropeptide expression is affected in a *ceh-34* mutant (irrespective of whether the effect is partial or total). See legend to Figure 3 for more information on features of circuit diagram.

We first examined the expression of the phylogenetically conserved Thyrotropin-releasing hormone (TRH) signaling axis that is important in stimulated gastrointestinal motility in vertebrates (Abot et al., 2018). *C. elegans* homologs of either the Thyrotropin-releasing hormone (TRH-1) or its receptor (TRHR-1) are expressed in the pharyngeal nervous system (Hunt-Newbury et al., 2007; Van Sinay et al., 2017), a notion we confirmed and extended with CRISPR/Cas9-engineered reporter alleles, showing that *trh-1* is expressed in the MI, M4 and M5 neurons and *trhr-1* in the M1, M2, M3 and I5 neurons (**Fig.4**, **Fig.5**). We find that the expression of the *trh-1* and *trhr-1* reporter alleles in these pharyngeal neurons is strongly affected in *ceh-34* mutants (**Fig.4**, **Fig.5**).

In addition to this deeply conserved neuropeptidergic system, we also tested the expression of a cohort of neuropeptides from the FMRFamides (*flp-2, flp-4, flp-5, flp-15, flp-21, flp-28*) and other miscellaneous neuropeptides (*nlp-3, nlp-8, nlp-13*). The expression of these nine neuropeptides is neuron-type specific, but in aggregate they cover the entire pharyngeal nervous system, often with unique cell-type specific combinations (schematized in **Fig.5B**). We find that the expression of all of these nine neuropeptides, analyzed with either reporter transgenes or CRISPR/Cas9 genome-engineered reporter alleles, is affected in *ceh-34* mutants (**Fig.5A**). We schematize these results again in the context of a pharyngeal circuit diagram to illustrate the breadth of effects of *ceh-34* on neuropeptide expression (**Fig.5B**). Together with our analysis of classic neurotransmitters (ACh, Glu, serotonin) and receptors, we conclude that *ceh-34* is required to endow pharyngeal neurons with their neuron type-specific arsenal of signaling molecules and, hence, that *ceh-34* is a critical specifier of several key aspects of pharyngeal neuron identity.

### *ceh-34* is required for sensory receptor expression in the pharyngeal nervous system

We sought to extend our analysis of *ceh-34* mutants by examining the expression of other molecular features of pharyngeal neurons. Like neurons in the vertebrate enteric nervous system, many of the pharyngeal neurons are likely sensory neurons that perceive sensory information to modulate peristaltic movements of the alimentary tract (Cook et al., 2020). While the sensory apparatus of pharyngeal neurons is not well understood, there are several candidate sensory receptors expressed in pharyngeal neurons. A gustatory receptor family member, *gur-3*, a possible light receptor (Bhatla and Horvitz, 2015), is expressed in two pharyngeal neuron classes and its expression is lost in *ceh-34* mutants (**Fig.6A**). Pharyngeal neurons also express the two sole members of the ionotropic sensory receptor family (Croset et al., 2010), encoded by *glr-7* and *glr-8* in *C. elegans* (Brockie et al., 2001; Hobert, 2013). We examined expression of *glr-7* expression, normally observed in six pharyngeal neuron classes (I2, I3, I6, M2, M3 and NSM), in *ceh-34* mutants and found its expression to be completely lost (**Fig.6A**). Finally we analyzed the expression of *str-97*, a putative chemosensory receptor of the GPCR family which we found to be expressed in several pharyngeal neurons (Vidal et al., 2018) (**Fig.6A**). We found that *ceh-34* is required for proper *str-97* expression (**Fig.6A**).

**Figure 6:**
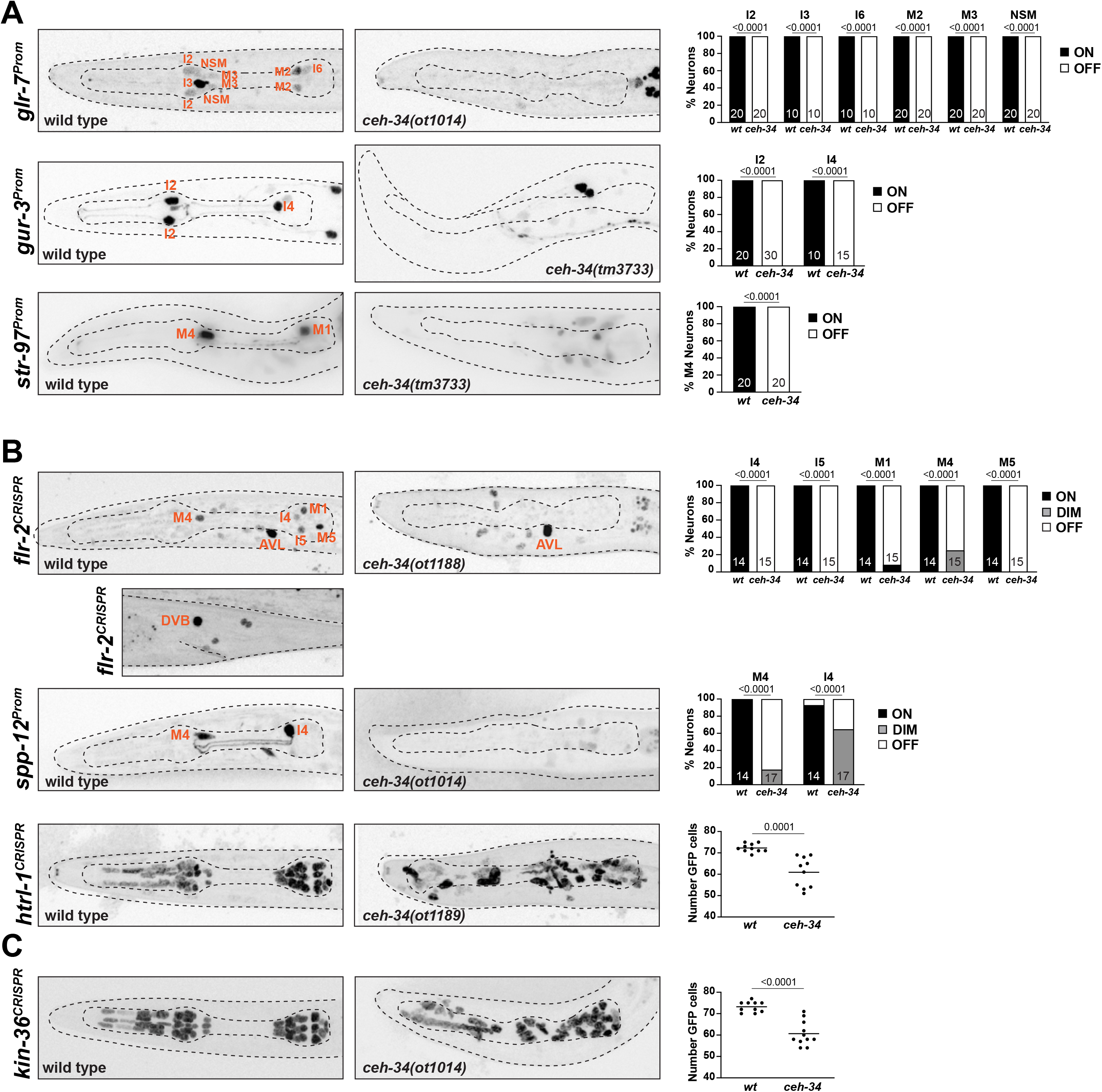
*ceh-34* affects other identity features of pharyngeal neurons. **(A)** Representative pictures and quantification showing sensory receptor expression loss in *ceh-34* mutants. Reporter genes used are *glr-7::gfp (otIs809), gur-3::gfp (nIs780)* and *str-97::gfp (otIs716)*. *str-97::gfp* is expressed in M1 in adult animals, but also in M4 in first larval stage animals. Expression in M4 is lost in *ceh-34* mutant, but expression in M1 could not be reliably scored. Statistical analysis was performed using Fisher’s exact test. N is indicated within each bar and represents number of neurons scored. **(B)** Representative pictures and quantification showing effect of *ceh-34* on antimicrobial defense genes. Reporter genes used are *flr-2 (syb4861), spp-12 (otIs868)* and *htrl-1 (syb4895)*. Statistical analysis was performed using Fisher’s exact test, Chi-Square test or unpaired t-test. N is indicated within each bar and represents number of neurons scored. **(C)** Representative pictures and quantification showing effect of *ceh-34* on pan-pharyngeal genes. Reporter gene is the CRISPR/Cas9-engineered reporter allele *kin-36 (syb4677).* Statistical analysis was performed using unpaired t-test.

### *ceh-34* is required for the expression of antimicrobial defense machinery

One deeply conserved feature of the gut and its associated nervous system is their engagement in antimicrobial defense, either directly through the release of antimicrobial peptides or through the employment of signaling systems that activate the immune system (Klimovich and Bosch, 2018; Muniz et al., 2012). Similar defense strategies operate in *C. elegans* (Dierking et al., 2016). Aside from the epithelial cells of the intestines, the pharyngeal nervous system appears to play a direct role in these microbial control mechanisms, as inferred by the pharyngeal neuron expression of specific proteins implicated in antimicrobial defense. For example, pharyngeal neurons express a hormone, FLR-2, homologous to glycoprotein hormone alpha subunit, that signals to the intestine to orchestrate antimicrobial defense (Oishi et al., 2009). Pharyngeal neurons also secrete pore-forming polypeptides that directly kill bacteria, such as the SPP-12 protein (Hoeckendorf et al., 2012), as well as a defensin-type antimicrobial peptide, ABF-2 and fungal-induced peptides (FIPR proteins)(Kato et al., 2002; Taylor et al., 2021). Our scRNA transcriptome analysis (Taylor et al., 2021) revealed pharyngeal neuron expression of another saposin-related secreted protein, which we named *htrl-1* (see Methods). We visualized expression of these signaling molecules, using a promoter fusion for *spp-12* (Hoeckendorf et al., 2012) and CRISPR/Cas9-engineered reporter alleles for *flr-2* and *htrl-1* (**Fig.6B**). We find that the *spp-12* reporter is expressed in I4 and M4 neurons, the *flr-2::SL2::gfp::h2b* reporter allele is expressed in 10 of the 14 pharyngeal neuron classes and the *htrl-1::SL2::gfp::h2b* reporter allele is expressed in all pharyngeal neurons (and in all other pharyngeal cells, but nowhere outside the pharynx)(**Fig.6B**). A notable feature of the extrapharyngeal expression of the *flr-2* reporter allele is expression in the AVL and DVB neurons (**Fig.6B**), the only extrapharyngeal neurons of the *C. elegans* nervous system that innervate gut tissue (not the foregut, but the midgut)(White et al., 1986). Crossing these reporters into a *ceh-34* mutant background, we found that expression of all three genes (*spp-12, flr-2, htrl-1*) is severely reduced and/or eliminated in pharyngeal neurons (**Fig.6B**).

### *ceh-34* is required for the expression of pan-pharyngeal nervous system genes

In addition to investigating the *ceh-34*-dependence of genes that fall into specific functional categories, we also sought to capitalize on the recently released single cell transcriptome profiling of the entire *C. elegans* nervous system that included the entire pharyngeal nervous system (Taylor et al., 2021). This analysis had shown that the molecular signatures of pharyngeal neurons are more similar to each other than to other neurons in the nervous system (Taylor et al., 2021)(**Fig.1D**). This pattern is driven, in part, by:

a. the pan-pharyngeal neuron, but pharynx-exclusive expression of *ceh-34,* as well as its transcriptional cofactor, Eyes absent/*eya-1* (described in more detail below),
b. pharynx restricted expression of the serotonin receptor *ser-7*, and several neuropeptides (e.g. *tkr-1*), all *ceh-34* targets, as described above,
c. a number of previously entirely uncharacterized genes with very broad, if not pan-pharyngeal nervous system expression (but no expression in non-pharyngeal neurons), including the above-mentioned saposin-related *htrl-1* gene, small cell surface proteins (e.g. C54E4.4) and a novel receptor tyrosine kinase with distant homology to the insulin receptor (Panther domain PTHR24416:SF525a), which we named *kin-36* (**Fig.6 – Supplement 1**). To confirm this expression pattern, we tagged the *kin-36* locus with a *gfp::H2B::SL2* cassette at it’s 5’ end, using CRISPR/Cas9 genome engineering. We found that *gfp::kin-36* indeed displays pan-pharyngeal neuron expression; expression is also observed in all other pharyngeal cell types, but no cell types outside the pharynx (**Fig.6C**). This pattern is strikingly similar to that of the saposin-related *htrl-1* reporter allele (**Fig.6B**), making *htrl-1* and *kin-36* the only known genes whose expression is so exclusively coupled to all pharyngeal cell types, but no other cell type. Crossing *gfp::kin-36* into *ceh-34* mutants, we observe what appears to be selective loss of *kin-36* expression from many, albeit not all pharyngeal neurons, a similar effect to what we observed for *htrl-1* (**Fig.6B, C**).

We conclude that *ceh-34* is required for the adoption of a broad palette of individual molecular features of all pharyngeal neurons (**Table 1**), consistent with a role as a terminal selector of neuronal identity of all pharyngeal neurons. Like other terminal selectors, *ceh-34* only affects neuron-type specific features, but not features that are expressed by all neurons throughout the nervous system.

**Table 1:**
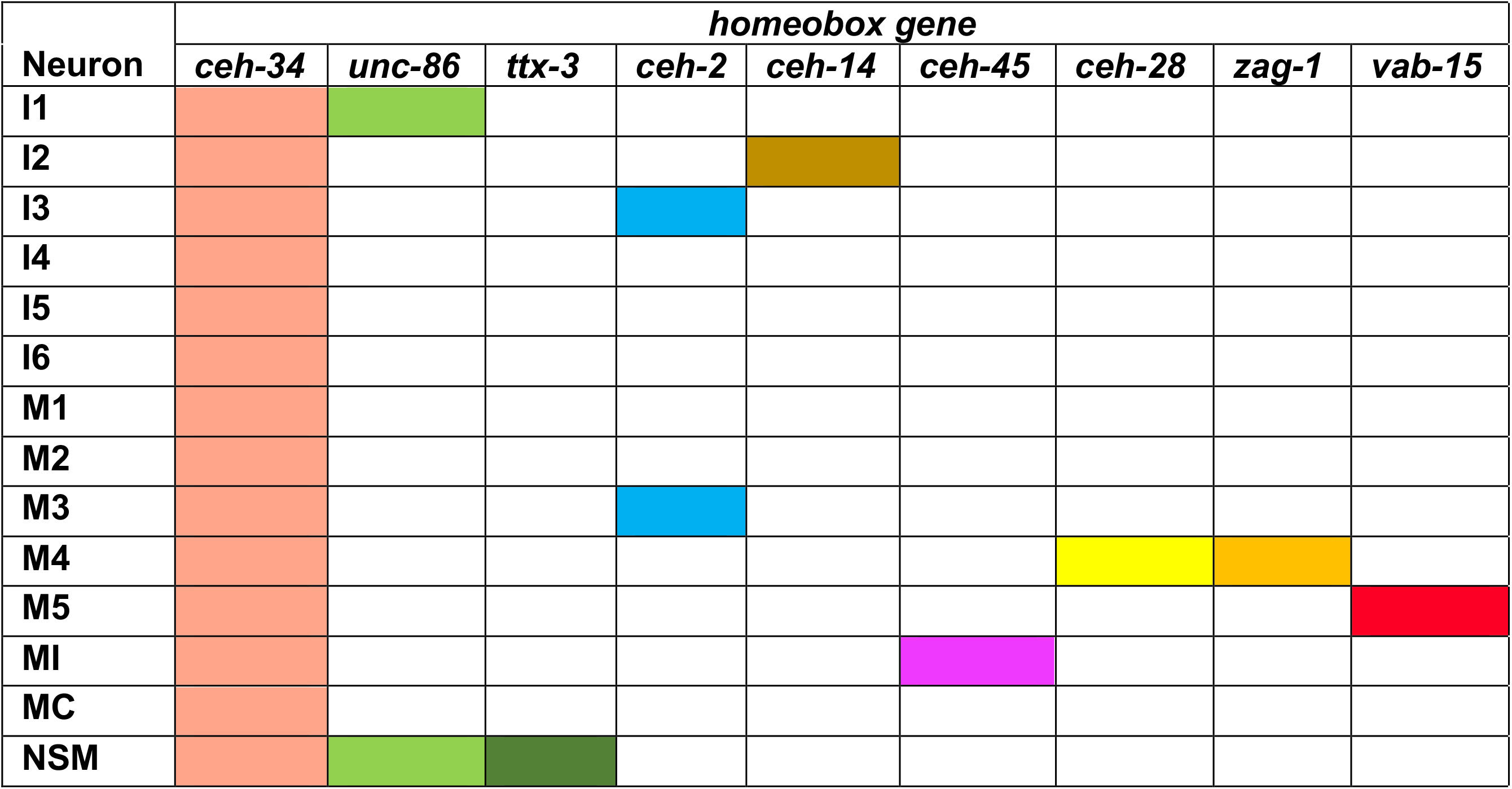
Tabular summary of homeobox codes involved in pharyngeal neuron identity specification.

### *ceh-34* is continuously required to maintain the differentiated state

The effect of *ceh-34* on terminal marker expression and its continuous expression throughout the life of all pharyngeal neurons suggests that, like other terminal selectors in the non-pharyngeal nervous system, *ceh-34* may not only initiate but also maintain the terminally differentiated state. To test this possibility, we generated a conditional *ceh-34* allele that allowed us to deplete CEH-34 protein postdevelopmentally. To this end, we inserted an auxin-inducible degron (Zhang et al., 2015) into the *ceh-34* locus using CRISPR/Cas9 genome engineering. Together with a ubiquitously expressed TIR1 ubiquitin ligase that recognizes the degron (*eft-3* driver; *ieSi*57)(Zhang et al., 2015), this approach allows for temporal depletion of CEH-34 protein through addition of auxin to the plate. (**Fig.7A**) Constitutive auxin exposure does not phenocopy the larval arrest phenotype of *ceh-34* null mutants, consistent with the persistence of very low level of CEH-34::mNG::AID (**Fig.7**). Nevertheless, postembryonic auxin addition resulted in downregulation of two tested markers for the differentiated state of different pharyngeal neuron classes, *eat-4/VgluT* and *unc-17/VAChT* (**Fig.7**). We conclude that *ceh-34* is required to maintain differentiated features of pharyngeal neurons and therefore fulfills another key criterion to classify as a terminal selector of pharyngeal neuron identity.

**Figure 7:**
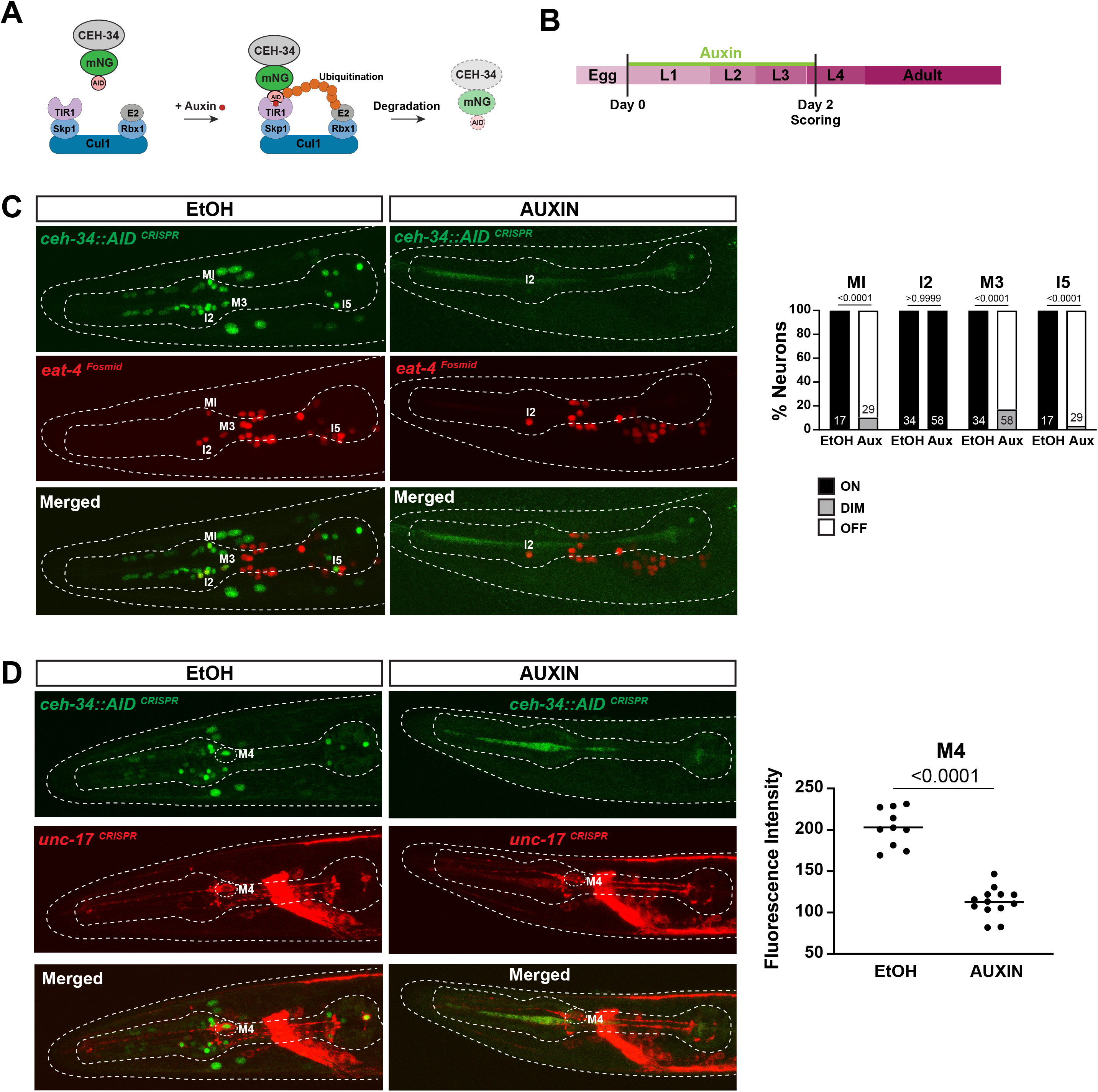
*ceh-34* is continuously required for identity maintenance. **(A)** Schematic of the AID system (Zhang et al., 2015). Skp1, Cul1, Rbx1 and E2 are phylogenetically conserved components of the E3 ligase complex. TIR1 is a plant-specific substrate-recognizing subunit of the E3 ligase complex. In the presence of auxin, the enzyme TIR1 binds to the AID fused to a protein of interest, leading to ubiquitination and proteasomal degradation of the targeted protein. **(B)** Schematic depicting the auxin treatment. Auxin was added to synchronized populations of worms at the L1 stage. Expression of molecular identity markers was analyzed in animals after 2 days on ethanol as vehicle control or 2 days on auxin. Worms were expressing TIR1 ubiquitously under the *eft-3* promoter (ieSi57). The *ceh-34* locus was tagged with *mNG::AID (ot903)*. **(C)** *ceh-34* is required for maintenance of *eat-4* expression (*otIs518*). Representative pictures and quantification are shown. Statistical analysis was performed using Chi-Square test. N is indicated within each bar and represents number of neurons scored. **(D)** *ceh-34* is required for maintenance of *unc-17* expression. Reporter gene used is a non-nuclear *unc-17* reporter allele (*ot907*), in which the *unc-17* locus is fused to a reporter gene. Representative pictures and quantification are shown. Statistical analysis was performed using unpaired t-test.

### Pharyngeal nervous system architecture is severely disorganized in *ceh-34* mutants

We further extended our analysis of *ceh-34* function by analyzing the anatomy of pharyngeal neuron circuitry in *ceh-34* mutants. This analysis is particularly important in light of the observation that, both in *C. elegans* (Berghoff et al., 2021; Pereira et al., 2015) and in vertebrates (Brunet and Pattyn, 2002), several instances have been described in which synaptically interconnected, but otherwise distinct neurons express the same transcription factor. Such observation suggests that these transcription factors may have a role in assembling neurons into functional circuitry. *ceh-34* represents a particularly extreme version of this scenario, because all *ceh-34(+)* neurons are heavily synaptically interconnected and only make a single robust synaptic contact to the rest of the *C. elegans* nervous system (**Fig.1C**)(Cook et al., 2020). To assess whether *ceh-34* not only specifies terminal molecular properties of a neuron, but also organizes overall circuit architecture, we examined pharyngeal nervous system architecture using fluorescent reporter constructs. Such analysis is complicated by the fact that genes that are selectively expressed in *ceh-34(+)* neurons, and hence could serve as drivers for a fluorescent reporter, are turned off in *ceh-34* null mutant animals, thereby preventing an easy visualization of individual axonal tracts or synaptic contacts. However, we found that the *ceh-34* promoter itself is still expressed until the first larval stage in *ceh-34* null mutants, when these animals arrest development. This transgene (*otIs762)* reveals that although their identity is not properly specified, as described above, pharyngeal neurons retain the capability to grow neuronal projections in *ceh-34* mutants; however, axonal tracts are severely disorganized (**Fig.8A**).

**Figure 8:**
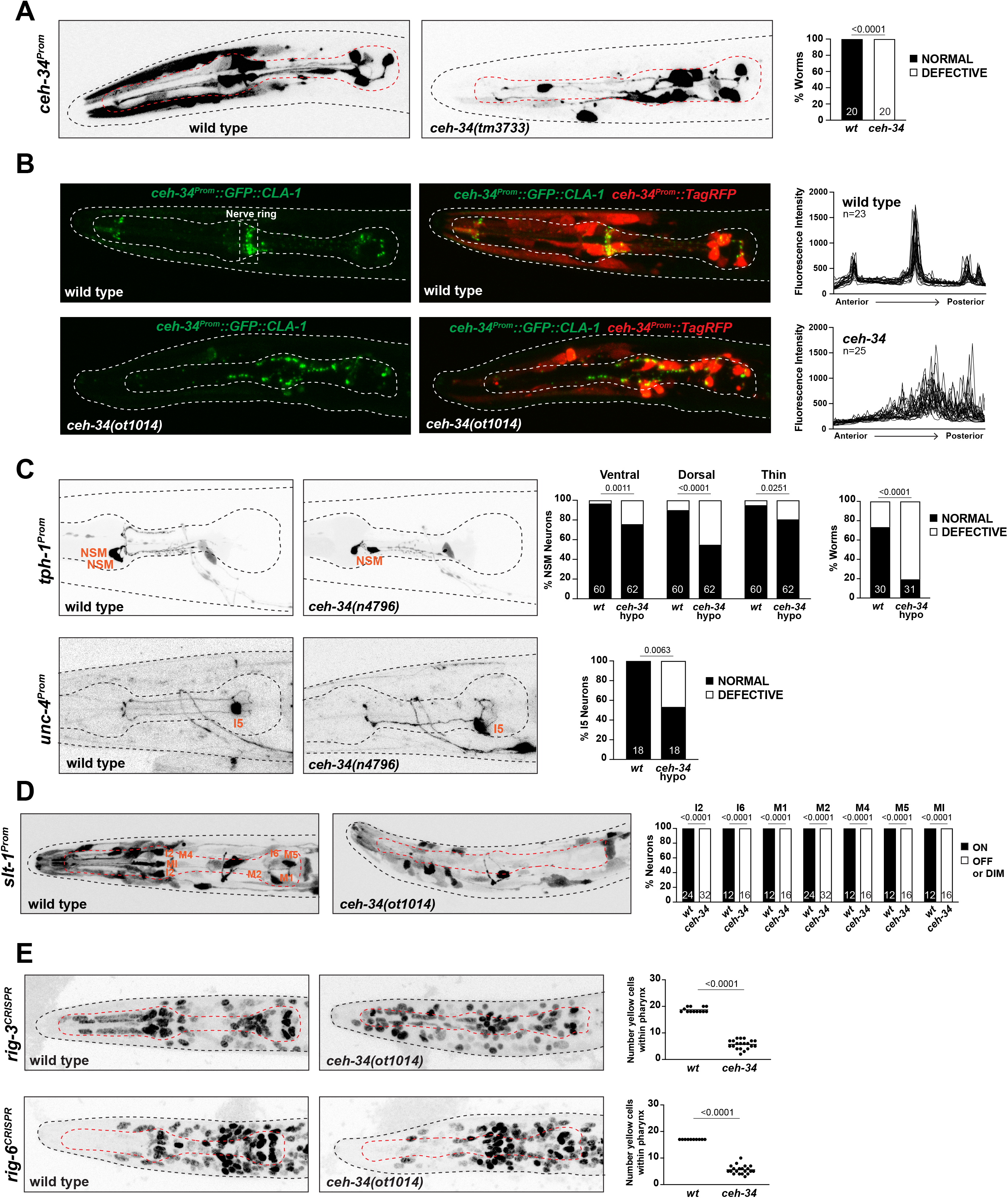
*ceh-34* affects the assembly of pharyngeal circuitry. **(A)** *ceh-34* null mutants have very disorganized axonal projections. Representative pictures and quantification are shown. Reporter gene is *ceh-34* (*otIs762*). Statistical analysis was performed using Fisher’s exact test. N is indicated within each bar and represents number of worms. **(B)** *ceh-34* null mutants show disorganized pharyngeal nerve ring presynaptic specializations as visualized with CLA-1 puncta. Representative pictures are shown. Quantification (right panels) shows gfp intensity profiles along the anterior posterior axis. Reporter gene is *otIs785*. **(C)** *ceh-34(n4796)* hypomorph mutants show axonal defects in NSM (top panel) and I5 (bottom panel). Representative pictures and quantification are shown. For NSM the ventral, dorsal and thin projection (not visible in picture) were scored separately (graph on the left) and then data was pulled together to indicate the percentage of worms showing any defect (graph on the right). Reporter genes used are *tph-1* (*zdIs13*) and *unc-4* (*otEx7503*). Statistical analysis was performed using Fisher’s exact test. N is indicated within each bar and represents number of neurons or number of worms. **(D)** *ceh-34* affects expression of the axon guidance cue *slt-1* (*kyIs174*). Representative pictures and quantification are shown. Statistical analysis was performed using Fisher’s exact test. N is indicated within each bar and represents number of neurons scored. **(E)** *ceh-34* affects expression of CRISPR/Cas9 engineered reporter alleles for *rig-3 (syb4763)* and *rig-6 (syb4729)*, two Ig superfamily members*. rig-3* and *rig-6* are expressed in almost all pharyngeal neurons plus many other cells within and outside the pharynx. Worms were scored with a red panneuronal marker (*otIs355*) or a red *ceh-34promoter* fusion (*stIs10447*) in the background to facilitate scoring. Number of yellow cells were counted within the pharynx. Statistical analysis was performed using unpaired t-test.

We also expressed the cytoplasmically localized TagRFP reporter together with a synaptically localized, GFP-tagged CLA-1/Clarinet protein (a synaptic active zone marker)(Xuan et al., 2017) under control of the *ceh-34* promoter. In wild-type animals, this transgene (*otIs785)* reveals (a) the axonal tracts of the pharyngeal nervous system and (b) synaptic structures that are strongly enriched in the pharyngeal nerve ring (**Fig.8B**). In *ceh-34* null mutants, we observed not only a disruption of axonal tract anatomy, but a severe disorganization of synaptic clusters throughout the entire pharyngeal nervous system (**Fig.8B**).

Lastly, we made use of a mild *ceh-34* hypomorphic allele, *n4796* (Hirose et al., 2010). In *ceh-34(n4796)* animals, the expression of many molecular markers for individual pharyngeal neurons are not affected, allowing to visualize their morphology. Focusing on two neuron types, NSM and I5, we find that in both types, specific axons branches fail to form in *ceh-34(n4796)* animals (**Fig.8C**). We conclude that *ceh-34* is required to establish proper pharyngeal nervous system architecture.

### *ceh-34* affects the expression of molecules involved in proper wiring of the pharyngeal nervous system

To further explore how *ceh-34* may affect pharyngeal nervous system architecture, we considered the expression of molecules with potential or explicitly demonstrated roles in axon guidance and/or synapse formation. A genetic analysis of axon guidance and circuit formation has only been conducted in a small number of pharyngeal neurons, mainly the M2 and NSM neuron classes (Pilon, 2008). In both neuron classes, the SLT-1 axon guidance cue, the *C. elegans* orthologue of Slit, has been found to be required for proper M2 and NSM axon guidance (Axang et al., 2008; Rauthan et al., 2007). We extended this phenotypic characterization, finding that other pharyngeal neurons also display axon pathfinding defects in *slt-1* mutants (**Fig.8 – Supplement 1**). To examine potential links between *slt-1* and *ceh-34,* we made use of a promoter::gfp fusion that captures the entire upstream intergenic region of the *slt-1* locus (Hao et al., 2001) and which shows selective expression in seven pharyngeal neuron classes (I2, I6, M1, M2, M4, M5 and MI) at the first larval stage. We find that *slt-1* expression is essentially abrogated in *ceh-34* mutants (**Fig.8D**).

Effects of *ceh-34* on molecules potentially involved in circuit formation are not restricted to *slt-1*. Two Ig superfamily members, *rig-3* and *rig-6* (the sole *C. elegans* ortholog of contactin) have previously been implicated in axon outgrowth and synapse function in the *C. elegans* nervous ssytem (Babu et al., 2011; Bhardwaj et al., 2020; Katidou et al., 2013; Kim and Emmons, 2017) and promoter fusion transgenes have indicated their expression in pharyngeal neurons (Schwarz et al., 2009). We used CRISPR/Cas9 to tag both loci with *gfp* and found that both genes are broadly expressed in many pharyngeal neurons (**Fig.8E**). *ceh-34* affects the pharyngeal neuron expression of both of *rig-3* and *rig-6* reporter alleles (**Fig.8E**).

### The Six homeodomain cofactor, Eyes absent/Eya, cooperates with *ceh-34*

To gain further insights into how CEH-34 patterns the identity of a wide array of distinct pharyngeal neuron types, we considered the involvement of cell-type specific cofactors. As a first step, we considered the EYes Absent/EYA protein, a phylogenetically conserved transcriptional co-activator of specific subsets of Six homeodomain proteins (Ohto et al., 1999; Patrick et al., 2013; Tadjuidje and Hegde, 2013). The *C. elegans* ortholog of Eyes absent, called *eya-1* (Furuya et al., 2005), also directly physically interacts with CEH-34 protein (Amin et al., 2009; Hirose et al., 2010). We first examined *eya-1* expression in pharyngeal neurons using a genomic fragment that contains the entire, *gfp-*tagged *eya-1* locus (Furuya et al., 2005). We observed expression in all pharyngeal neurons throughout all larval and adult stages, a precise phenocopy of the *ceh-34* expression pattern (**Fig.9A**), an expression pattern also corroborated by our recent scRNA analysis (**Fig.6 – Supplement 1**)(Taylor et al., 2021). Moreover, we found that *eya-1* expression requires *ceh-34* function, suggesting that *ceh-34* acts in a feedforward configuration to induce its own transcriptional cofactor (**Fig.9B**).

**Figure 9:**
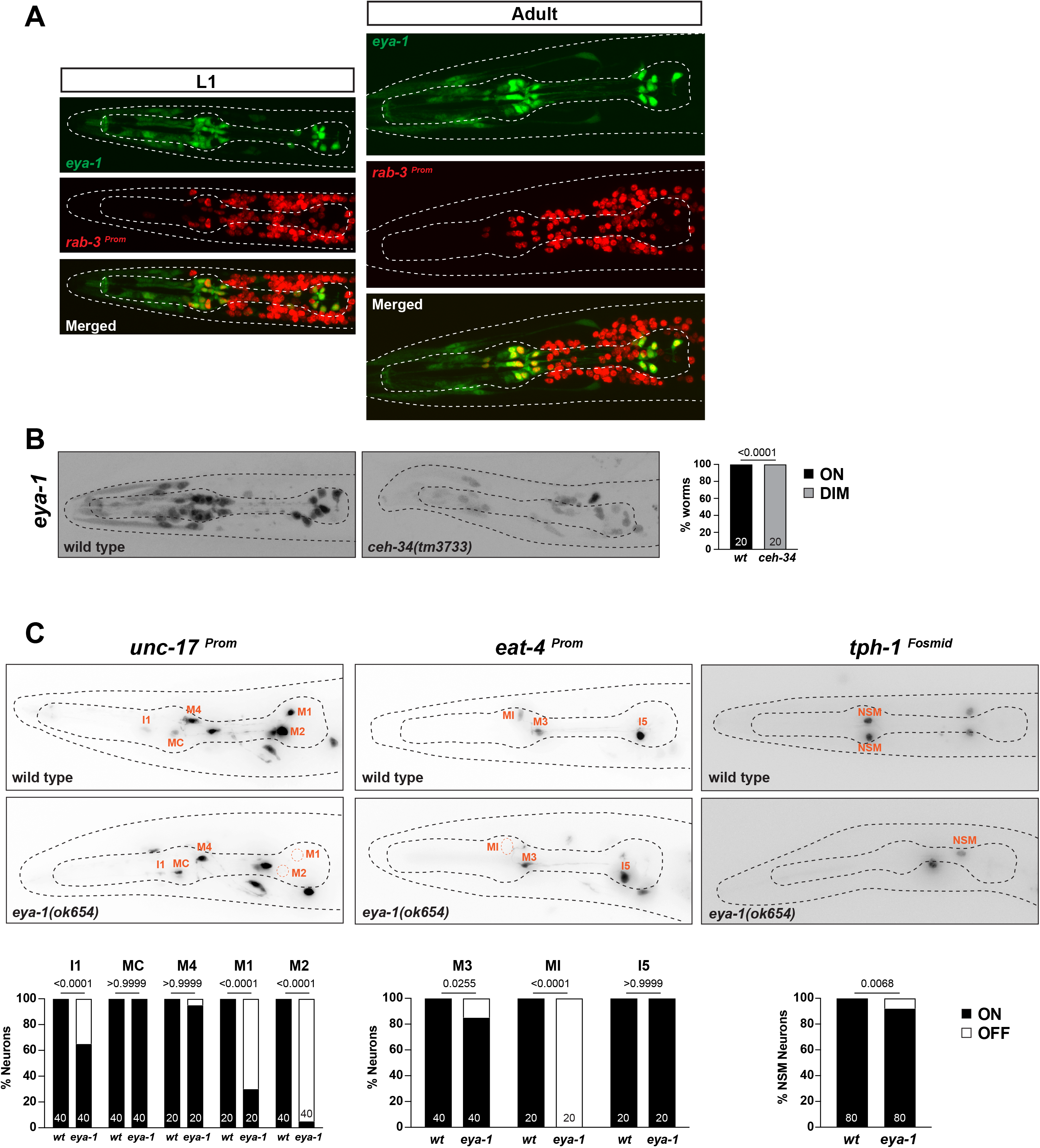
*ceh-34* cooperates with the *eya-1* general co-factor. **(A)** *eya-1* is expressed in all pharyngeal neurons throughout the life of the worm. Images of L1 and adult worms showing co-localization of *eya-1* expression (*nIs352*) with the panneuronal gene *rab-3* (*otIs355*) in pharyngeal neurons. **(B)** *eya-1* expression is regulated by *ceh-34.* Representative pictures and quantification are shown. Reporter gene is *eya-1* (*nIs352*). Statistical analysis was performed using Fisher’s exact test. N is indicated within each bar and represents number of worms scored. **(C)** *eya-1* mutant shows defects in pharyngeal neuron neurotransmitter identity. Representative images and quantification are shown for *unc-17 (otIs661), eat-4 (otIs487)* and *tph-1 (otIs517)*. Statistical analysis was performed using Fisher’s exact test. N is indicated within each bar and represents number of neurons scored.

We analyzed the function of *eya-1* in the context of pharyngeal neuron specification. Animals that carry a deletion of a part of the *eya-1* locus, *ok654,* display pharyngeal neuron specification defects, albeit much milder than those observed in *ceh-34* null mutant animals (**Fig.9C**). The larval arrest phenotype of *ceh-34* null mutants is also more penetrant than that of *eya-1* mutants, which are homozygous viable (Furuya et al., 2005). Since it is not clear whether the *eya-1* allele is a null allele, it remains unclear whether *ceh-34* may be able to partly function without *eya-1*.

In other organisms, Dachshund proteins are components of Sine oculis/Eya complexes in several cellular contexts (Hanson, 2001). However, the sole *C. elegans* ortholog of Dachshund, *dac-1,* does not cooperate with CEH-34/EYA-1 in the context of pharynx development. First, *dac-1* is not expressed in pharyngeal neurons (Colosimo et al., 2004; Taylor et al., 2021) and, second, *dac-1* null mutants do not display the larval arrest phenotype characteristic of *ceh-34* and *eya-1* mutants (Colosimo et al., 2004).

### *ceh-34* cooperates with a multitude of other homeobox genes to specify distinct pharyngeal neuron types

How does *ceh-34* activate distinct genes in different pharyngeal neuron types? One obvious possibility is that *ceh-34* cooperates with neuron type-specific cofactors in neuron-type specific terminal selector complexes to drive specific fates. As candidates for such cofactors, we considered homeobox genes, for two reasons: (1) like any other neuron in the *C. elegans* nervous system, each individual pharyngeal neuron expresses a unique combination of homeobox genes, in addition to pan-pharyngeal *ceh-34* (Reilly et al., 2020); (2) previous studies had already implicated a few homeobox genes in controlling some select functional or molecular aspects of individual pharyngeal neurons (Aspock et al., 2003; Feng and Hope, 2013; Morck et al., 2004; Ramakrishnan and Okkema, 2014; Ray et al., 2008; Zhang et al., 2014). For example, the homeobox gene *ceh-2,* the *C. elegans* ortholog of vertebrate EMX and Drosophila Ems, is required for proper function of the M3 neuron, but effects of *ceh-2* on molecular aspects of M3 neuron differentiation had not been reported (Aspock et al., 2003). We therefore set out to analyze homeobox gene function throughout the pharyngeal nervous system and to examine interaction of *ceh-34* with other homeobox genes.

#### NSM neurons

To ask whether distinct pharyngeal neuron type-specific homeobox genes cooperate with *ceh-34* in distinct neuronal cell types throughout the pharyngeal nervous system, we made use of a hypomorphic *ceh-34* allele, *n4796* (Hirose et al., 2010). Unlike the *ceh-34* null allele, which displays highly if not completely penetrant effects on all NSM molecular markers (**Fig.3-6**), the *n4796* allele displays only subtle if any neuronal differentiation defects on its own (**Fig.10A**). However, when combined a mutant allele of *unc-86,* a POU homeobox gene that affects many but not all NSM marker genes (Zhang et al., 2014) strong synergistic differentiation defects of NSM are observed (**Fig.10A**). This genetic interaction mirrors the synergistic effects of *unc-86* and the LIM homeobox gene *ttx-3*, another regulator of NSM differentiation (Zhang et al., 2014).

**Figure 10:**
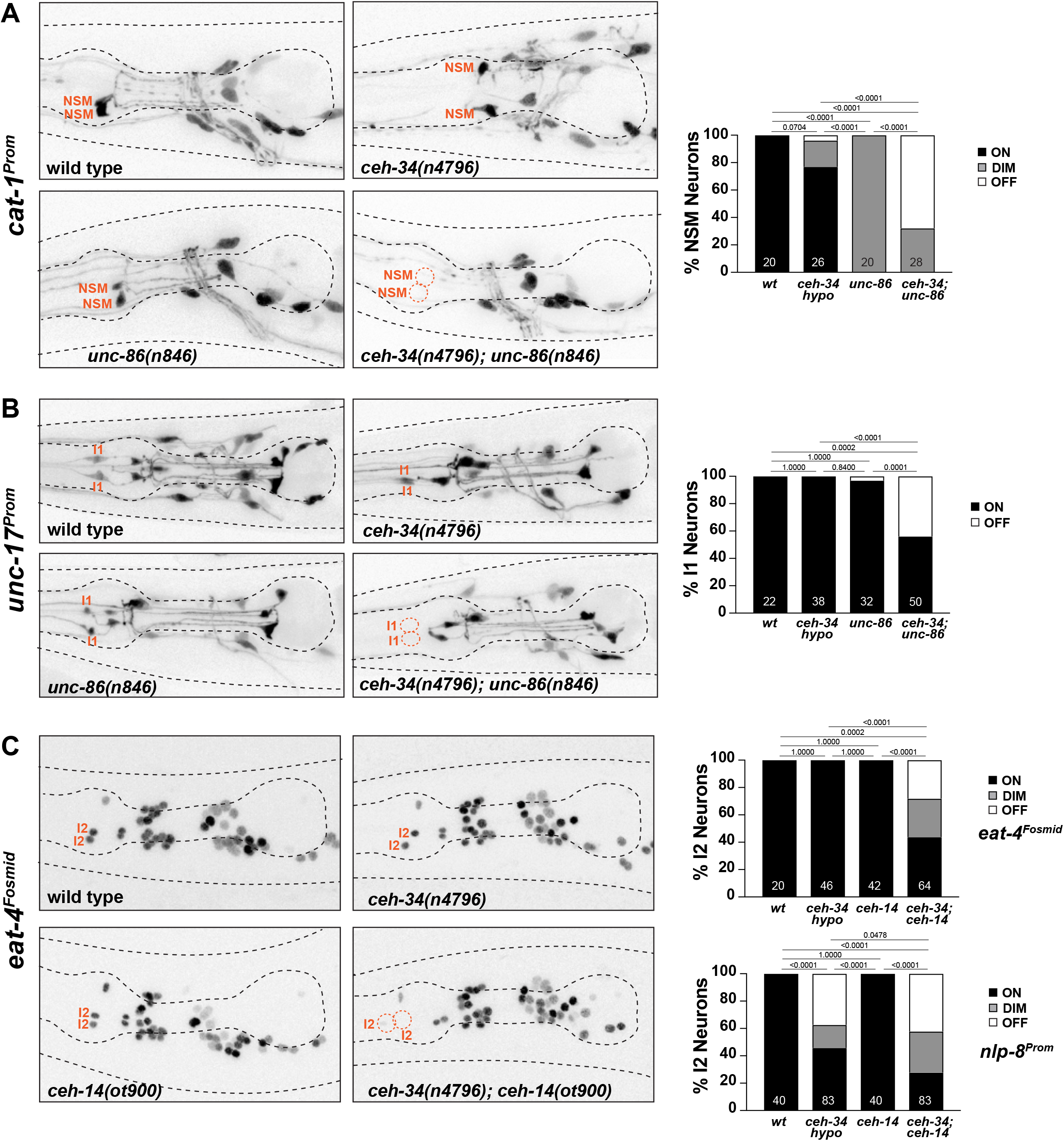
*ceh-34* cooperates with LIM and POU-type homeobox genes to specify distinct pharyngeal neuron types. **(A)** *unc-86* and *ceh-34* show synergistic effects in NSM differentiation. Representative images and quantification are shown. Reporter gene used is *cat-1* (*otIs224*). Statistical analysis was performed using Fisher’s exact test. P-values were adjusted with the Holm-Sidak correction for multiple comparisons. N is indicated within each bar and represents number of neurons scored. **(B)** *unc-86* and *ceh-34* show synergistic effects in I1 neuron differentiation. Representative images and quantification are shown. Reporter gene used is *unc-17* (*otIs661*). Statistical analysis was performed using Fisher’s exact test. P-values were adjusted with the Holm-Sidak correction for multiple comparisons. N is indicated within each bar and represents number of neurons scored. **(C)** *ceh-14* and *ceh-34* show synergistic effects in I2 neuron differentiation. Representative images and quantification are shown for *eat-4* (*otIs518*). Bottom graph shows quantification for *nlp-8* (*otIs711*). Statistical analysis was performed using Fisher’s exact test or Chi-Square test . P-values were adjusted with the Holm-Sidak correction for multiple comparisons. N is indicated within each bar and represents number of neurons scored.

#### I1 Sneurons

A similar genetic interaction between *ceh-34* and *unc-86* is observed in the cholinergic I1 neuron pair, the only other pharyngeal neuron class that also co-expresses *ceh-34* and *unc-86* (Baumeister et al., 1996; Serrano-Saiz et al., 2018), but which does not express *ttx-3*. While cholinergic identity, visualized via *unc-17/VAChT* expression, is not affected in *unc-86(n846)* single mutants, a combination of the *ceh-34(n4796)* hypomorphic allele with the *unc-86(n846)* mutation results in I1 losing its cholinergic identity (**Fig.10B**).

#### I2 neurons

Synergistic interactions are also observed in the glutamatergic I2 neuron pair. Like the I1 neurons, no identity regulators were previously known for this neuron class. Our homeobox gene mapping project (Reilly et al., 2020) showed that the I2 neuron expresses the LIM homeobox gene *ceh-14,* the *C. elegans* ortholog of vertebrate Lhx3/4. No other pharyngeal neuron expresses *ceh-14*. While *ceh-14* single null mutants show no effect on *eat-4/VGLUT* expression (the marker of glutamatergic identity of the I2 neurons), in combination with the *ceh-34(n4796)* hypomorphic allele a strong synergistic effect on glutamatergic identity acquisition is observed in I2 (**Fig.10C**). Another molecular marker for I2 identity, the neuropeptide *nlp-8*, is also synergistically regulated by *ceh-34* and *ceh-14* (**Fig.10C**).

#### I3 neuron

The previously unstudied cholinergic I3 neuron class expresses, in addition to *ceh-34*, the *C. elegans* ortholog of Empty spiracles/EMX, *ceh-2* (Aspock et al., 2003; Reilly et al., 2020). The *ceh-2* null mutant allele, *ch4,* alone show a reduction of expression of the *unc-17/VAChT* reporter allele (**Fig.11A**). In combination with the *ceh-34(n4796)* hypomorphic allele *unc-17/VAChT* expression and, hence, cholinergic identity of I3, is eliminated (**Fig.11A**).

#### M3 neuron

Apart from expression in I3, the EMX ortholog *ceh-2* is also expressed in the glutamatergic M3 neurons and is required for M3 function (Aspock et al., 2003), but molecular correlates for this functional defect have not previously been identified. We find that in *ceh-2* single mutants, expression of the *eat-4/VGluT* identity marker is affected in M3 (**Fig.11B**).

#### M4 neuron

In the cholinergic M4 neuron, the *ceh-28* and *zag-1* homeobox genes have previously been shown to each regulate subsets of M4 identity feature (Ramakrishnan and Okkema, 2014). Both *zag-1* and *ceh-28* affected *flp-2* expression, but only *zag-1*, and not *ceh-28,* was found to affect *ser-7* expression (Ramakrishnan and Okkema, 2014). In contrast, *ceh-28* but not *zag-*1 affected *flp-5* expression and neither *zag-1* nor *ceh-28* affected *unc-17* or *flp-21* expression (Ramakrishnan and Okkema, 2014). *ceh-34* null mutants show effects on the expression of all the tested *ceh-28* or *zag-1*-dependent (*ser-7, flp-2* and *flp-5*) or independent markers (*unc-17, flp-21*)(**Fig.3-5)**, indicating that *ceh-34* may collaborate with these homeobox genes to control distinct subsets of M4 differentiation markers.

#### *M5* neuron

The cholinergic M5 neuron expresses, in addition to *ceh-34*, the Msh/Msx ortholog *vab-15* (Reilly et al., 2020). Since only a hypomorphic allele of *vab-15* was previously available (Du and Chalfie, 2001), we generated a molecular null allele, *ot1136*, through CRISPR/Cas9 genome engineering (**Fig.11 – Supplement 1**). We found a complete loss of expression of the cholinergic marker *unc-17/VAChT* expression, as well as of the neuropeptidergic marker, *trh-1,* in *vab-15(ot1136)* animals (**Fig.11C**). Complete losses of marker expression are not an indicator of failure of this neuron to be generated, since crossing a *ceh-34* marker into *vab-15* mutants revealed the presence and normal *ceh-34* expression of M5 (**Fig.11C**).

**Figure 11:**
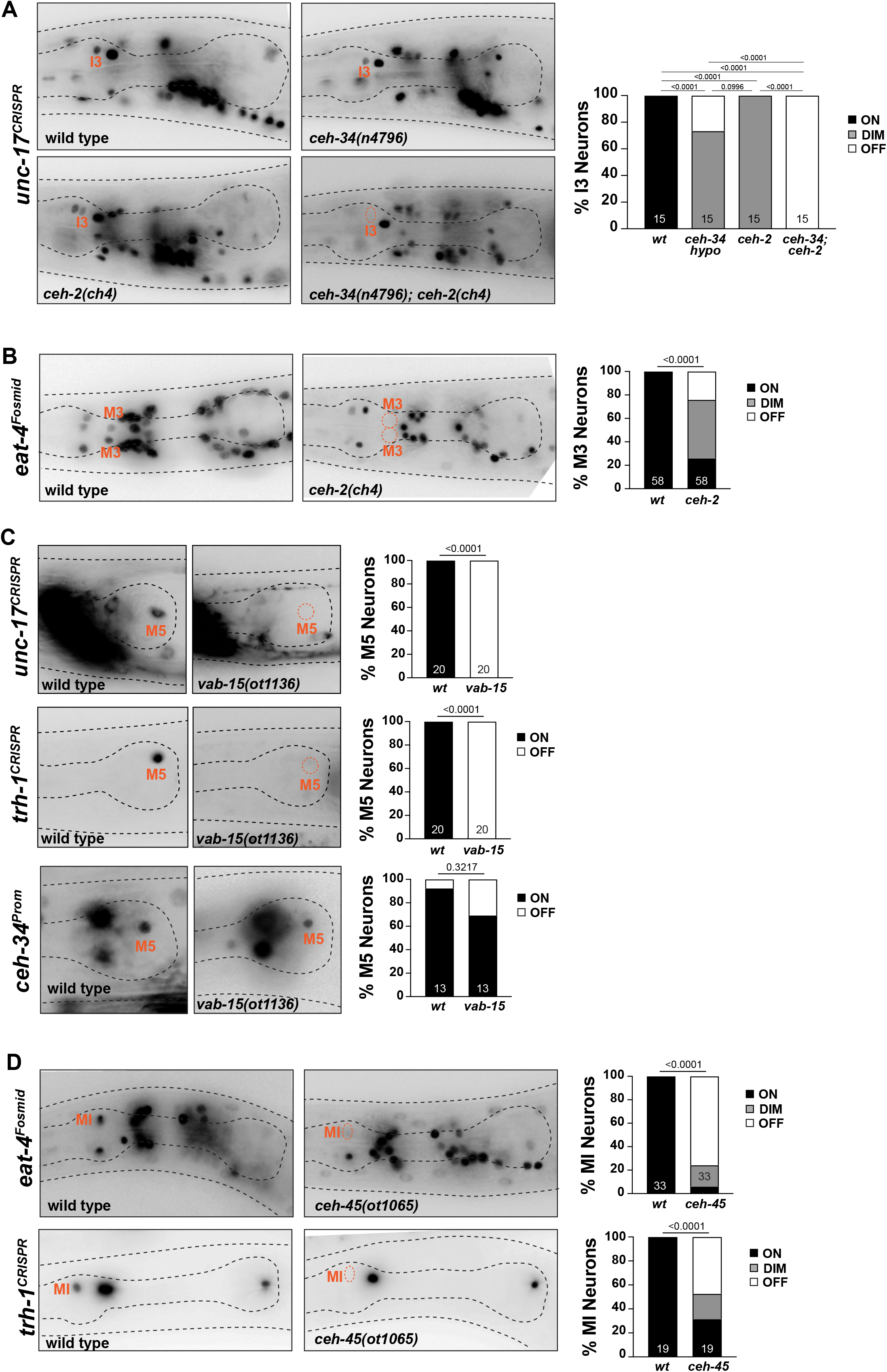
Other homeobox genes involved in specifying distinct pharyngeal neuron types. **(A)** *ceh-2* and *ceh-34* show synergistic effects in I3 neuron differentiation. Representative images and quantification are shown. Reporter gene used is *unc-17* (*syb4491*). Statistical analysis was performed using Fisher’s exact test. P-values were adjusted with the Holm-Sidak correction for multiple comparisons. N is indicated within each bar and represents number of neurons scored. **(B)** *ceh-2* affects M3 neuron differentiation. Representative images and quantification are shown. Reporter gene is *eat-4* (*otIs388*). Statistical analysis was performed using Chi-Square test. N is indicated within each bar and represents number of neurons scored. **(C)** *vab-15* affects M5 neuron differentiation. Representative images and quantification are shown. Reporter alleles used are *unc-17* (*ot907*), *trh-1* (*syb4421*) and *ceh-34* (*stIs10447*). Statistical analysis was performed using Fisher’s exact test. N is indicated within each bar and represents number of neurons scored. **(D)** *ceh-45* affects MI neuron differentiation. Representative images and quantification are shown. Reporter genes used are *eat-4* (*otIs388*) and *trh-1* (*syb4421*). Statistical analysis was performed using Fisher’s exact test. N is indicated within each bar and represents number of neurons scored.

#### MI neuron

In our previous genome-wide analysis of homeobox gene expression (Reilly et al., 2020), we had shown that the glutamatergic MI neuron expresses the sole worm ortholog of the Goosecoid homeobox gene, *ceh-45.* Embryonically, *ceh-45* is expressed in multiple pharyngeal tissues (Ma et al., 2021), but its expression resolves to exclusive expression in the MI, as well as the I1 neuron (Reilly et al., 2020). *ceh-45* had not previously been functionally uncharacterized. We examined *ceh-45* function by generating a null allele using CRISPR/Cas9 genome engineering, *ot1065* (**Fig.11 – Supplement 1**). Mirroring the *ceh-34* defects, we found that glutamatergic identity specification of MI (as assessed by *eat-4/VGLUT* expression) is strongly affected in *ceh-45* null mutant animals (**Fig.11D**). Similarly, expression of the neuropeptide *trh-1* is also affected in MI (**Fig.11D**).

In conclusion, eight homeobox genes appear to collaborate with *ceh-34* in eight of the 14 pharyngeal neuron classes to specify their proper identity (**Table 1**). Since the remaining six classes also express specific combinations of homeobox genes (Reilly et al., 2020), we anticipate that future analysis will likely reveal homeobox codes throughout the entire pharyngeal nervous system.

## DISCUSSION

We have identified here common, overarching themes of enteric nervous system differentiation in the nematode *C. elegans.* A number of previous studies have identified transcription factors involved in regulating specific differentiation aspects of a small subset of pharyngeal neurons (Aspock et al., 2003; Feng and Hope, 2013; Morck et al., 2004; Ramakrishnan and Okkema, 2014; Rauthan et al., 2007; Ray et al., 2008; Zhang et al., 2014), yet no common theme emerged from these studies. We have shown here that a Sine oculis ortholog, *ceh-34,* operates together with its phylogenetically conserved transcriptional coactivator *eya-1*, to selectively specify the terminal differentiation program of all pharyngeal neurons. The *ceh-34/eya-1* complex appears to act as a terminal selector of pharyngeal neuron identity, as inferred from its requirement to initiate terminal neuronal differentiation programs in all pharyngeal neurons (without overall loss of panneuronal identity, a unifying trait of all terminal selectors), as well as its continuous role in maintaining the differentiated state (another defining trait of terminal selectors). The aspects of the differentiation program that we consider here, and found to be under control of *ceh-34,* include anatomical (axon outgrowth and synapse formation), molecular and presumptive functional features. Molecular and functional features affected by *ceh-34* range from neuron-neuron communication to presumptive sensory functions to the intriguing function of enteric neurons as potential regulators of microbial colonization. There is good reason to believe that CEH-34 controls these diverse phenotypic identity features in a direct manner, i.e. it may not act through intermediary factors. CEH-34 is among the many *C. elegans* transcription factors whose binding sites have been determined *in vitro* through protein binding microarrays (Narasimhan et al., 2015) and, using a phylogenetic footprinting pipeline, we find that these motifs are significantly enriched in the single cell transcriptome of most pharyngeal neuron classes (Glenwinkel et al., 2021). This is in accordance with CEH-34 being a shared terminal selector component of all pharyngeal neuron classes.

Within the nervous system, the selectivity of *ceh-34* expression in all pharyngeal neurons is remarkable – based on extensive gene expression pattern analysis, including recent scRNA data, there is no other transcription factor that so selectively and comprehensively defines all enteric, but no non-enteric neuronal cell types. We found that the key determinant of this expression is the organ selector PHA-4 (Gaudet and Mango, 2002; Horner et al., 1998; Kalb et al., 1998; Mango et al., 1994), which is expressed earlier in development to act both as a pioneer factor (Hsu et al., 2015) and to induce the expression of a number of different terminal selectors for different tissue types within the foregut – the *ceh-34* gene for all neurons (this paper), the bHLH transcription factor *hlh-6* for pharyngeal gland cells (Smit et al., 2008) and the Nk-type homeobox gene *ceh-22* for pharyngeal muscle identity (Vilimas et al., 2004). Given the continuous pharyngeal expression of PHA-4 throughout postembryonic life that we observe, it is conceivable that PHA-4 acts in a regulatory feedforward motif configuration, where it not only induces tissue-type terminal selectors (*ceh-34, hlh-6, ceh-22*), but then also collaborates with them to induce and maintain terminal differentiation batteries.

Our work indicates that CEH-34 is a shared component of neuron-type specific terminal selector complexes, such that CEH-34 interacts with a distinct set of at least eight homeodomain co-factors to impose unique features in distinct pharyngeal neuron types. This *in vitro-*determined binding sites for these homeodomain co-factors display, like the CEH-34 site, a phylogenetically conserved enrichment in the respective neuron type-specific gene batteries (Glenwinkel et al., 2021). For example, CEH-34 and UNC-86 binding sites are co-enriched in the I1 neuronal transcriptome, CEH-34 and CEH-14 binding sites in the I2 neuronal transcriptome, CEH-34 and CEH-2 in the I3 transcriptome, CEH-34 and CEH-45 in the MI transcriptome and VAB-15 and CEH-34 in the M5 transcriptome (Glenwinkel et al., 2021).

Taken together, our findings not only reveal a common terminal selector-based regulatory logic for how a self-contained, enteric nervous system acquires its terminal differentiated state. They also corroborate two themes that have emerged from recent studies, primarily in *C. elegans* (and also emerging in other systems):

1. homeobox genes – and more specifically, homeobox gene combinatorial codes – are prominently employed in neuron identity specification throughout all neurons of the nervous system, as exemplified here in the context of the enteric nervous system, the so-called “second brain” of animals (Gershon, 1998).
2. A number of identity-specifying terminal selectors, such as CEH-34, are expressed in synaptically connected neurons, suggesting they may specify the assembly of neurons into functional circuitry. It is presently unclear how prominent such a connectivity theme is. We observed a few cases in the non-pharyngeal nervous system (Berghoff et al., 2021; Pereira et al., 2015) and there are some striking examples in vertebrates (Brunet and Pattyn, 2002; Dauger et al., 2003; Ha and Dougherty, 2018; Ruiz-Reig et al., 2019; Sokolowski et al., 2015). This present study provides an extreme example of this – an entire set of synaptically connected neurons (the worm’s enteric nervous system) is specified by a single transcription factor (CEH-34), which apparently helps these neurons in being assembled into functional circuitry. In each pharyngeal neuron type, CEH-34 pairs up with different homeodomain proteins to diversify pharyngeal neurons into distinct identities. Following Dobzhansky’s dictum that “nothing in biology makes sense except in the light of evolution” (Dobzhansky, 1964) we speculate that the pharyngeal nervous system may have derived from a homogenous set of interconnected, identical neurons, all specified by *ceh-34*, which may have regulated a homophilic adhesion molecule that functionally linked these ancestral neurons. The more complex, present day circuitry may have evolved through the eventual partnering of CEH-34 protein with additional homeodomain proteins that diversified neuronal identities, connectivity and function in the pharyngeal nervous system.

Arguing for a conserved function of Six homeodomain factors in enteric nervous system differentiation is the observation that in flies, the Sine oculis paralog Optix is indeed expressed in the frontal ganglion (Seo et al., 1999), which constitutes the nervous system of insect foreguts (Hartenstein, 1997). In the context of studying Sine Oculis function in the *Drosophila* corpus cardiacum, it was also noted that the entire stomatogastric ganglion, i.e. the entire enteric nervous system (of which the frontal ganglion is a part) does not form in Sine oculis mutants (De Velasco et al., 2004). In sea urchin, pharyngeal neurons also express, and require for their proper development, the Sine oculis paralog Six3 (Wei et al., 2011). Other than an early report of Six2 expression in the mouse foregut region (Ohto et al., 1998), the expression and function of Sine oculis orthologs in vertebrate enteric nervous systems has not yet been examined.

Notably, another homeobox gene appears to have a critical and very broad function in vertebrate enteric nervous system development that is akin to the broadness of *ceh-34* function in *C. elegans*. The Prd-type homeobox gene Phox2b is expressed in enteric nervous system precursors and required early in development for generation of all enteric ganglia (Pattyn et al., 1999; Tiveron et al., 1996). While Phox2b is also continuously expressed throughout the adult enteric nervous system (Corpening et al., 2008; Drokhlyansky et al., 2020; Morarach et al., 2021), its function in terminal differentiation and perhaps even maintenance of enteric neurons identity remains to be examined, for example, via temporally controlled, post-developmental knock-out in juvenile or adult stage animals. If the analogy to *ceh-34* holds, the enteric neurons of such animals may lose their differentiated state. Remarkably, additional homeobox genes have recently been noted to show highly selective expression patterns within the vertebrate enteric nervous system, effectively discriminating distinct neuronal subtypes (Memic et al., 2018). One of them, the Meis ortholog Pbx3 has been confirmed to play an important role in the postmitotic specification of distinct enteric neuron types (Morarach et al., 2021). Hence, it appears that the overall logic of a pan-enteric homeobox gene, cooperating with cell type specific homeobox genes, may be conserved from worms to vertebrates.

Another evolutionary perspective of our findings considers the origins of the enteric nervous system, and maybe nervous systems as a whole. Based on a number of anatomical and functional features, it has been proposed that enteric nervous systems preceded, and then paralleled the existence of centralized nervous system of bilaterian animals (Furness and Stebbing, 2018; Gilbert, 2019; Klimovich and Bosch, 2018). This argument is bolstered by considering a number of features of the enteric, i.e. pharyngeal nervous system of *C. elegans:* (a) its polymodality (sensory + inter + motor neuron) of most pharyngeal neurons, (b) its innervation of what is essentially a single sheath of myoepithelial cells, a proposed feature of primitive nervous systems (Mackie, 1970), (c) its simple immune functions (also thought to be a feature of primitive neurons, e.g. in hydra)(Klimovich et al., 2020; Klimovich and Bosch, 2018) and (d) the relatively indiscriminate synaptic cross-innervation patterns among pharyngeal neurons (Cook et al., 2020). If pharyngeal neurons indeed resemble a more primitive, ancestral state of neurons, our observation that CEH-34 acts as a terminal selector in these neurons would point to the ancient nature of (a) a terminal selector-type logic of neuronal identity specification and (b) the deployment of a homeobox gene in such function. Sine Oculis homologs appear to be employed broadly in sensory neuron specification across animal phylogeny, even in the most basal metazoan (Jacobs et al., 2007). CEH-34/Sine Oculis may represent an ancestral determinant of neuronal cell types.

## MATERIALS AND METHODS

### Microscopy

Worms were anesthetized using 100 mM sodium azide (NaN_3_) and mounted on 5% agarose pads on glass slides. Z-stack images (each ∼0.7 micron thick) were acquired using a Zeiss confocal microscope (LSM880) or Zeiss compound microscope (Imager Z2) using the ZEN software. Maximum intensity projections of 2 to 30 slices were generated with the ImageJ software (Schindelin et al., 2012).

### Caenorhabditis elegans strains

Worms were grown at 20°C on nematode growth media (NGM) plates seeded with *E. coli* (OP50) bacteria as a food source. The wild-type strain used is Bristol N2. A complete list of strains used in this study can be found in **Supplemental File 1**.

#### Generation of deletion alleles

Mutant alleles for the *ceh-34, ceh-45 and vab-15* genes (schematized in **Fig.2A** and **Fig.11 – Supplement 1**) were generated by CRISPR/Cas9 genome engineering as described (Dokshin et al., 2018). A deletion of the full locus was generated using two crRNAs and an ssODN donor. Sequences are as follows:

##### ceh-34(ot1014)

crRNAs (cgacaagaggacgacgctct and ttattctaatggtcttgagg), ssODN (gcgacattcactgggggacgacaagaggacgacgccaagaccattagaataacttttaactatatttttg).

*ceh-34(ot1188)* and *ceh-34(ot1189)* were generated the same way as *ceh-34(ot1014)* and are molecularly identical. The difference is that *ot1188* was generated in the background of *flr-2(syb4861)* and *ot1189* was generated in the background of *htrl-1(syb4895)* because these loci are very closely linked.

##### ceh-45(ot1065)

crRNAs (taggccaccgatacaagcag and tccgccagagaccggtcggg), ssODN (aactgaaattcgaaattctaggccaccgatacaaggaccggtctctggcggattactgtagccgtttggg).

##### vab-15(ot1136)

crRNAs (ggtcaacacatctgcttata and ttgtgaaaagcgtaatactt), ssODN (agcgcgtggtgttatattggtcaacacatctgctttattacgcttttcacaatattttatggactaacca).

#### Generation of reporter knock-ins

The *ceh-34* locus was tagged with *mNG::3xFLAG::AID* to generate *ceh-34(ot903)*. The Auxin-Inducible Degron (AID) sequence was amplified and inserted into the pDD268 vector (*mNG::SEC::3xFLAG*) (Dickinson et al., 2015) to generate the plasmid pUA77 (*ccdB::mNG::SEC::3xFLAG::AID ccdB*)(Aghayeva et al., 2021). The construct contains a self-excising drug selection cassette (SEC) and was used for SEC-mediated CRISPR insertion of *mNG::3xFLAG::AID* right before the stop codon of *ceh-34* as described in (Dickinson et al., 2015). The guide RNA used targets the following sequence: ttattctaatggtcttgagg.

The *pha-4* locus was tagged with *gfp* at its 3’end to generate *pha-4(ot946)*, using Cas9 protein, tracrRNA, and crRNA from IDT, as previously described (Dokshin et al., 2018). One crRNA (attggagatttataggttgg) and an asymmetric double stranded *gfp-loxP-3xFLAG* cassette, amplified from a plasmid, were used to insert the fluorescent tag at the C-terminal.

Reporter alleles for *flp-5(syb4513), flp-28(syb3207), flr-2(syb4861), htrl-1(syb4895), ser-7(syb4502), rig-3(syb4763), rig-6(syb4729), trh-1(syb4421) and trhr-1(syb4453)* were generated by CRISPR/Cas9 to insert an *SL2::GFP::H2B* cassette at the C-terminus of the respective gene. For the *unc-17(syb4491)* allele *T2A::GFP::H2B* was inserted at the C-terminus. For the *kin-36(syb4677)* locus, a *GFP::HIS::SL2* sequence was inserted at the N-terminus. These strains were generated by Sunybiotech and are listed in **Supplemental File 1**.

#### Generation of transgenic reporter strains

To generate *otIs762(ceh-34prom::TagRFP)* a 3720bp PCR fragment containing the whole *ceh-34* intergenic region plus the first 2 exons and 2 introns was amplified from N2 genomic DNA and cloned into a TagRFP vector using Gibson Assembly (NEBuilder HiFi DNA Assembly Master Mix, Catalog # E2621L). The following primers were used: aatgaaataagcttgcatgcctgcaTGTTTATTTTCTATGTAATTTCTAATAAAGTCCC and cccggggatcctctagagtcgacctgcaCTGAAAGTTGAAATATAGAATTTTTAATTTTTTTTTTTTG. The resulting construct was injected as a simple extrachromosomal array (50ng/ul) into *pha-1(e2123)* animals, using a *pha-1* rescuing plasmid (pBX, 50ng/ul) as co-injection marker. A representative line was integrated into the genome with gamma irradiation and backcrossed 4 times.

To generate *otIs785(ceh-34prom::GFP::CLA-1)* a 3720bp PCR fragment containing the whole *ceh-34* intergenic region plus the first 2 exons and 2 introns was amplified from N2 genomic DNA and cloned into PK065 (kindly shared by Peri Kurshan) using Gibson Assembly (NEBuilder HiFi DNA Assembly Master Mix, Catalog # E2621L). The following primers were used: gattacgccaagcttgcatg*c*TGTTTATTTTCTATGTAATTTCTAATAAAGTC and gttcttctcctttactcatc*ccggg*CTGAAAGTTGAAATATAGAATTTTTAATTTTTTTTTTTTG. The resulting construct was injected at 7ng/ul together with *ceh-34prom::TagRFP* (50ng/ul) (see above) and *rol-6(su1006)* as a co-injection marker. A representative line was integrated into the genome with gamma irradiation and backcrossed 4 times.

### Choice of pharyngeal fate markers

Most fate markers used in this paper were previously described. For several of those, we used previous expression patterns (based on transgenic reporter fusions) as a guide to then generate reporter alleles by CRISPR/Cas9 genome engineering (e.g. *flp-5*, *ser-7*; all listed in previous sections). One case warrants specific emphasis: scRNA has shown that the T11F9.12 gene is expressed exclusively in most, if not all pharyngeal neurons, a notion we confirmed with a CRISPR/Cas9 genome engineered reporter allele. T11F9.12 encodes for a relatively large (736aa), secreted and nematode specific protein that contains a Pfam-annoted domain (Htrl domain; PF09612) that is, outside nematodes, only found in a bacterial protein HtrL. This bacterial protein currently has no assigned function but is found in a region of LPS core biosynthesis genes which are involved in bacterial immune defense (Bertani and Ruiz, 2018). Interestingly, hidden Markov model-based searches in the Panther database reveal a sequence pattern (PTHR21579) along the entire T11F9.12 protein that is otherwise only found in *C. elegans* saposin proteins, which are bona fide immune effector proteins (Banyai and Patthy, 1998; Hoeckendorf et al., 2012; Oishi et al., 2009). We named this protein HTRL-1.

### Temporally controlled CEH-34 protein degradation

We used conditional protein depletion with the auxin-inducible degradation system (Zhang et al., 2015). AID-tagged proteins are conditionally degraded when exposed to auxin in the presence of TIR1. To generate the experimental strain, the conditional allele *ceh-34(ot903[ceh-34::mNG::AID])* was crossed with *ieSi57[eft-3^prom^::tir1]*, which expresses TIR1 ubiquitously. The natural auxin indole-3-acetic acid (IAA) was purchased from Alfa Aesar (Ward Hill, MA) (#A10556) and dissolved in ethanol (EtOH) to prepare 400-mM stock solutions. NGM agar plates with fully grown OP50 bacterial lawn were coated with the auxin stock solution to a final concentration of 4 mM and allowed to dry for a few hours at room temperature. To induce protein degradation, worms of the experimental strain were transferred onto the auxin-coated plates and kept at 20°C. As a control, worms were transferred onto EtOH-coated plates instead. Auxin solutions, auxin-coated plates, and experimental plates were shielded from light.

## ACKNOWLEDGEMENTS

We thank Chi Chen for generating transgenic lines, Molly Reilly and Seth Taylor for discussion, Kelly Liu, Robert Horvitz, Peri Kurshan and Matthias Leippe for providing reagents, Steven Cook for providing illustrations and members of the Hobert lab for comments on the manuscript. This work was funded by NIH R21NS106843 and the Howard Hughes Medical Institute.

---

**Figure 2 – Supplement 1:**
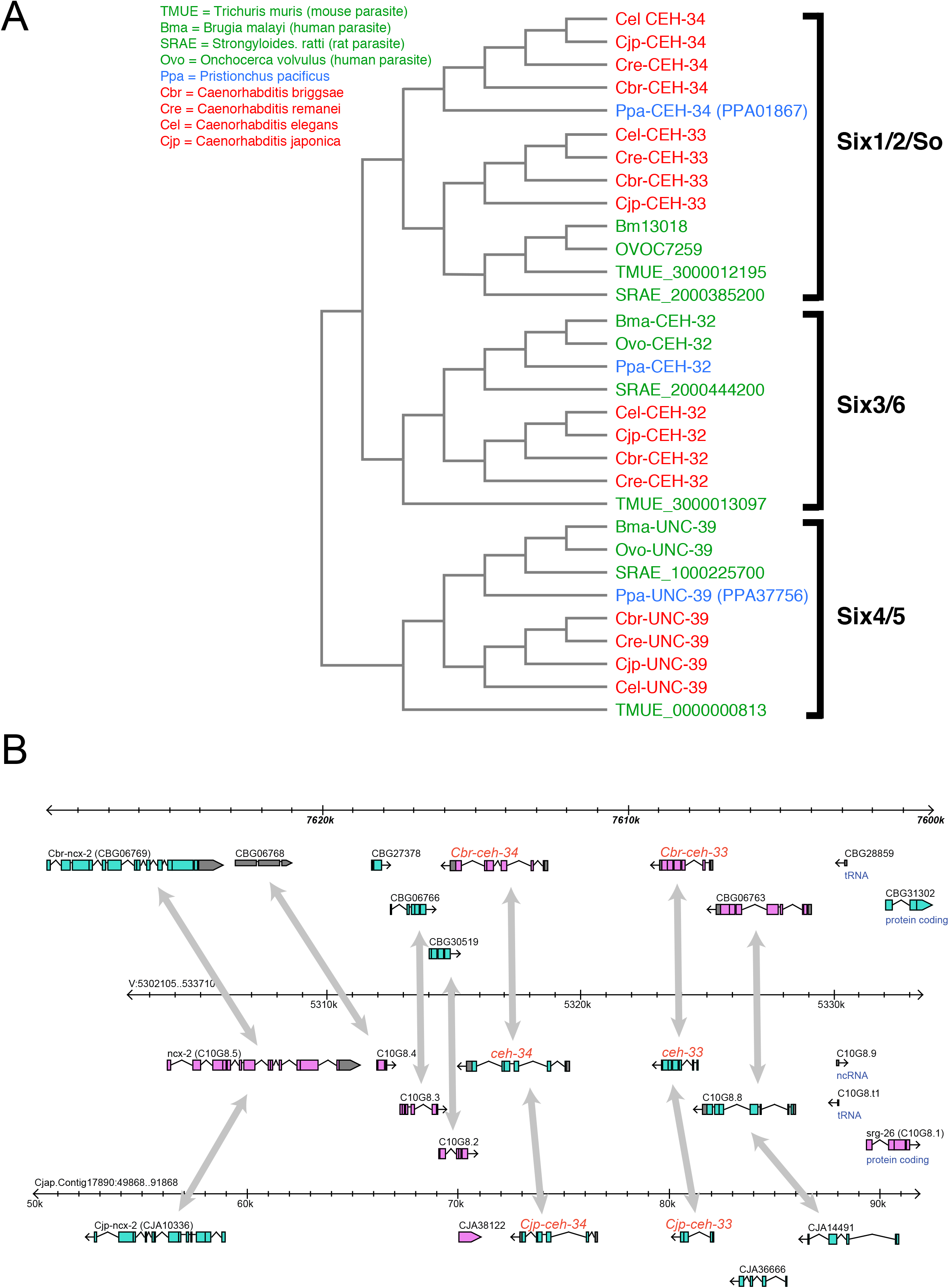
Phylogeny of *ceh-33/34* gene duplication in other nematodes. **(A)** Phylogeny based on homeodomains. **(B)** Synteny of the *ceh-33/ceh-34* locus across several *Caenorhabditis* species.

**Figure 2 – Supplement 2:**
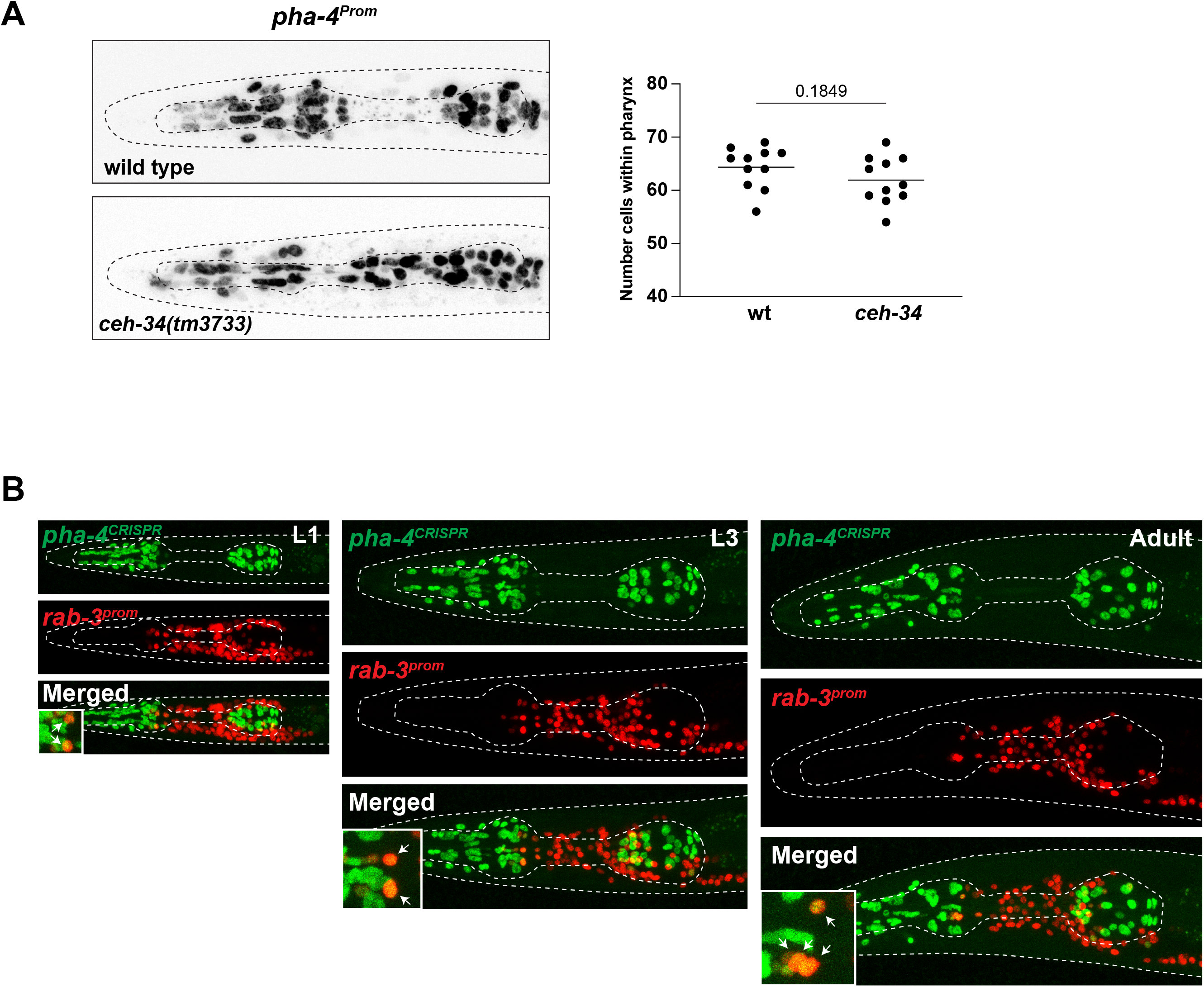
**(A)** *pha-4* expression (*stIs10077*) is not affected in *ceh-34* mutants. Statistical analysis was performed using unpaired t-test. **(B)** *pha-4* is continuously expressed in pharyngeal neurons. Expression of a CRISPR/Cas9-engineered *pha-4* reporter allele, “*pha-4^CRISPR)^“* (*ot946*) in green and a *rab-3^prom^* (*otIs355*) reporter in red in L1 (left), L3 (middle) and adult (right) animals. High magnifications of the anterior bulb provided in the insets at the bottom left corner of the merged images. Arrows point to pharyngeal neurons expressing both *pha-4* and *rab-3*.

**Figure 6 – Supplement 1:**
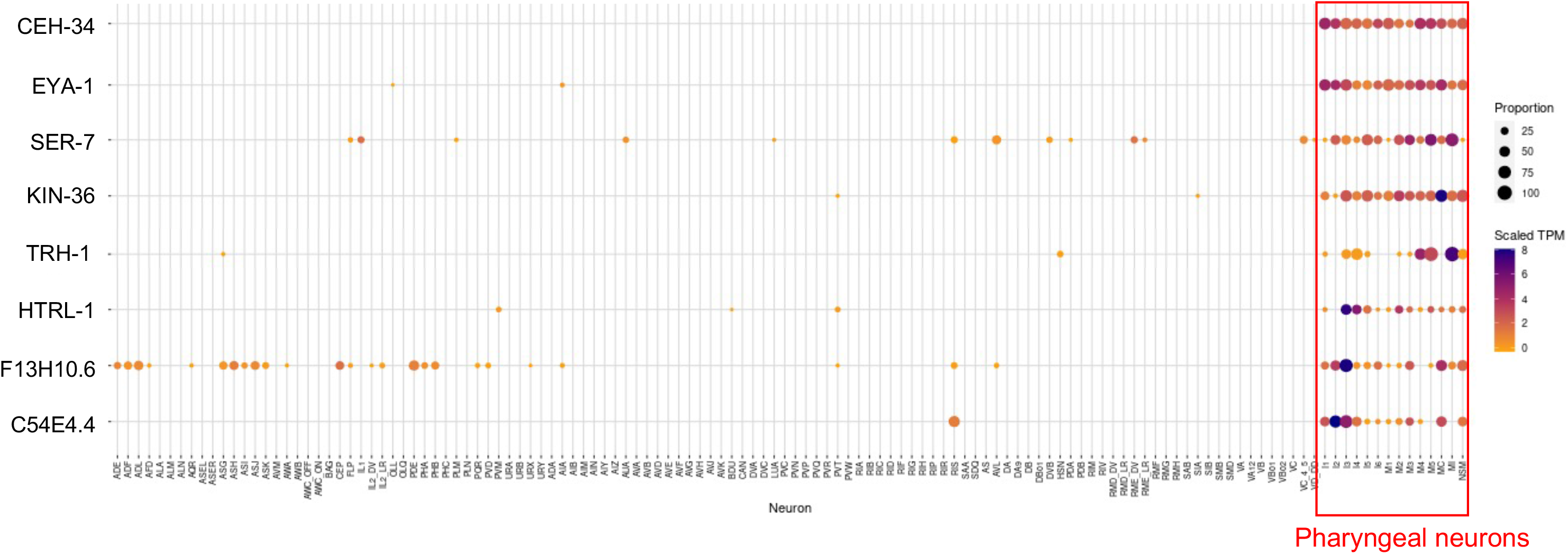
Single cell transcriptomic analysis. scRNA data was extracted from the Cengen App and displayed in a heatmap format, as previously described (Taylor et al., 2021). The F13H10.6 gene was included here as well because of its striking expression outside the pharyngeal nervous system, in what appear to be exclusively sensory neurons, in line with most if not all pharyngeal neurons also having sensory function. Conversely, previously identified pan-sensory genes (e.g. cilium related genes) do not show expression in pharyngeal neurons.

**Figure 8 – Supplement 1:**
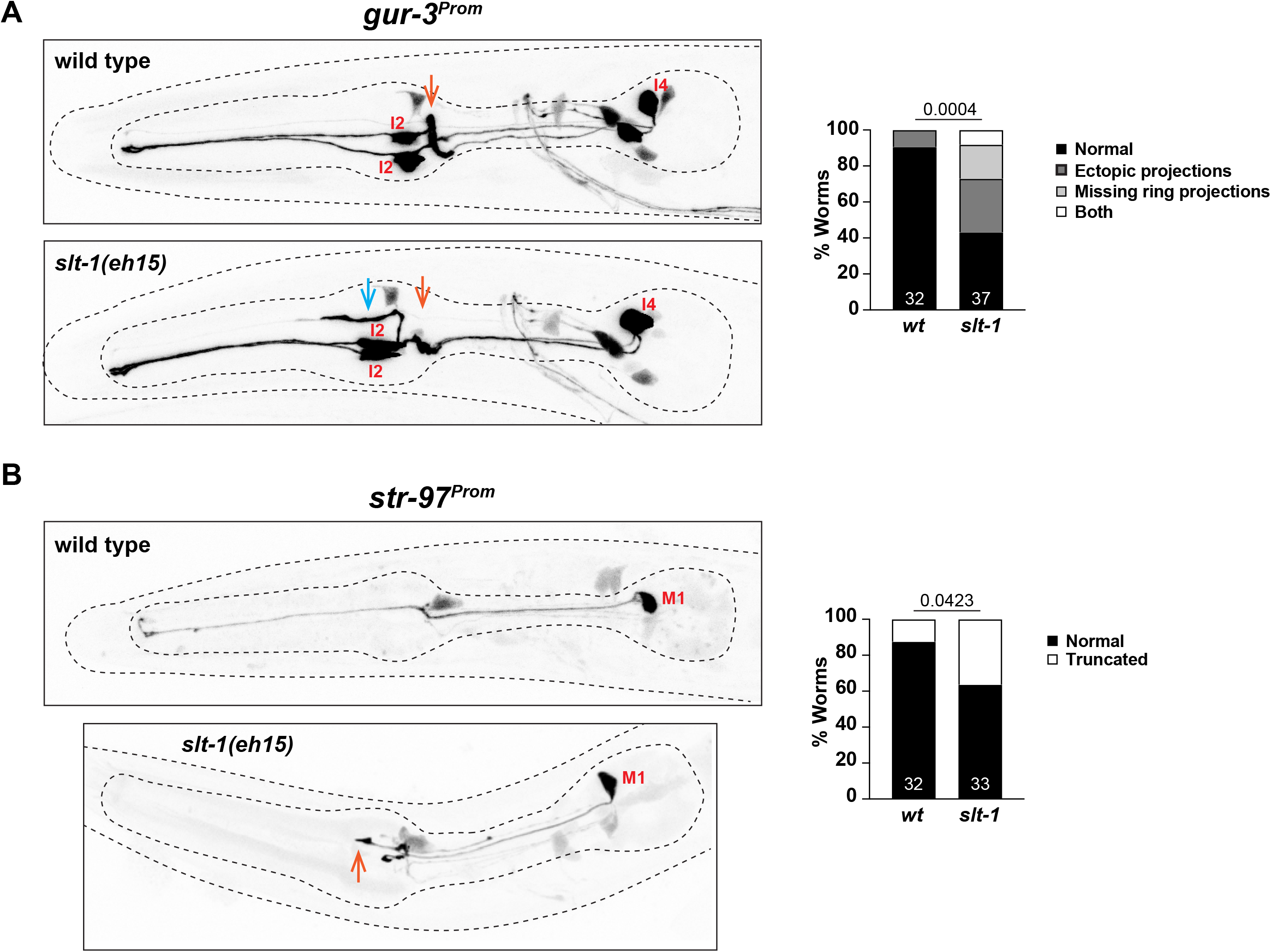
Axonal defects in *slt-1* mutants. Representative pictures and quantification are shown. **(A)** Effects of *slt-1* on the I2/I4 neurons. Orange arrows indicate position of pharyngeal nerve ring. Blue arrow indicates ectopic projection. Reporter gene is *gur-3* (*nIs780*). **(B)** Effects of *slt-1* on the M1 neuron. Orange arrow indicate shortened M1 process, which normally extends to the anterior end of the pharynx. Statistical analysis was performed using Chi-Square test. N is indicated within each bar and represents number of neurons scored.

**Figure 11 – Supplement 1:**
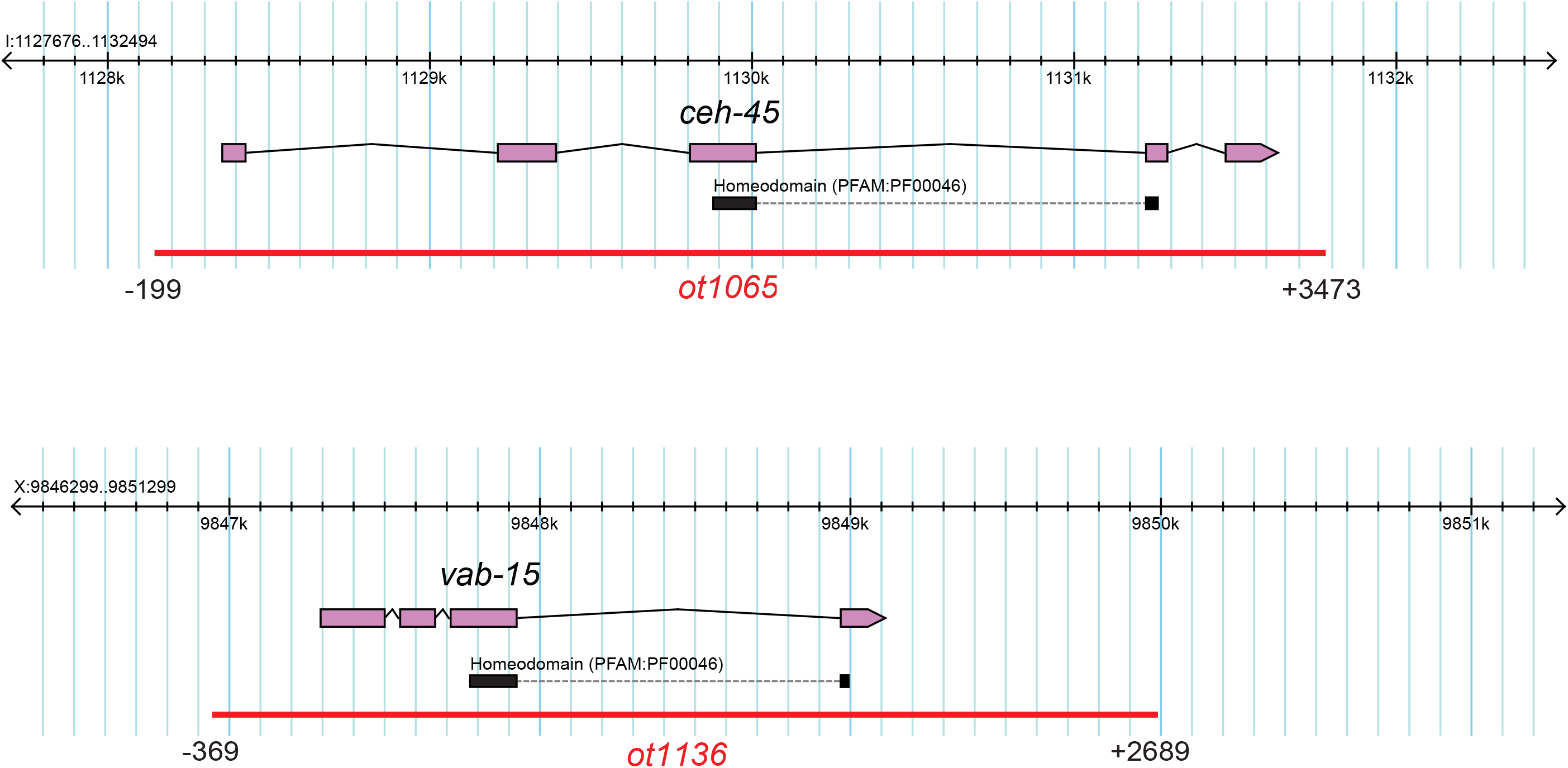
Homeobox mutant alleles generated in this study. Numbers indicating beginning and end of deletion are relative to the start codon of the gene of interest. See Methods for detail.

## Supplemental File 1: Strain List

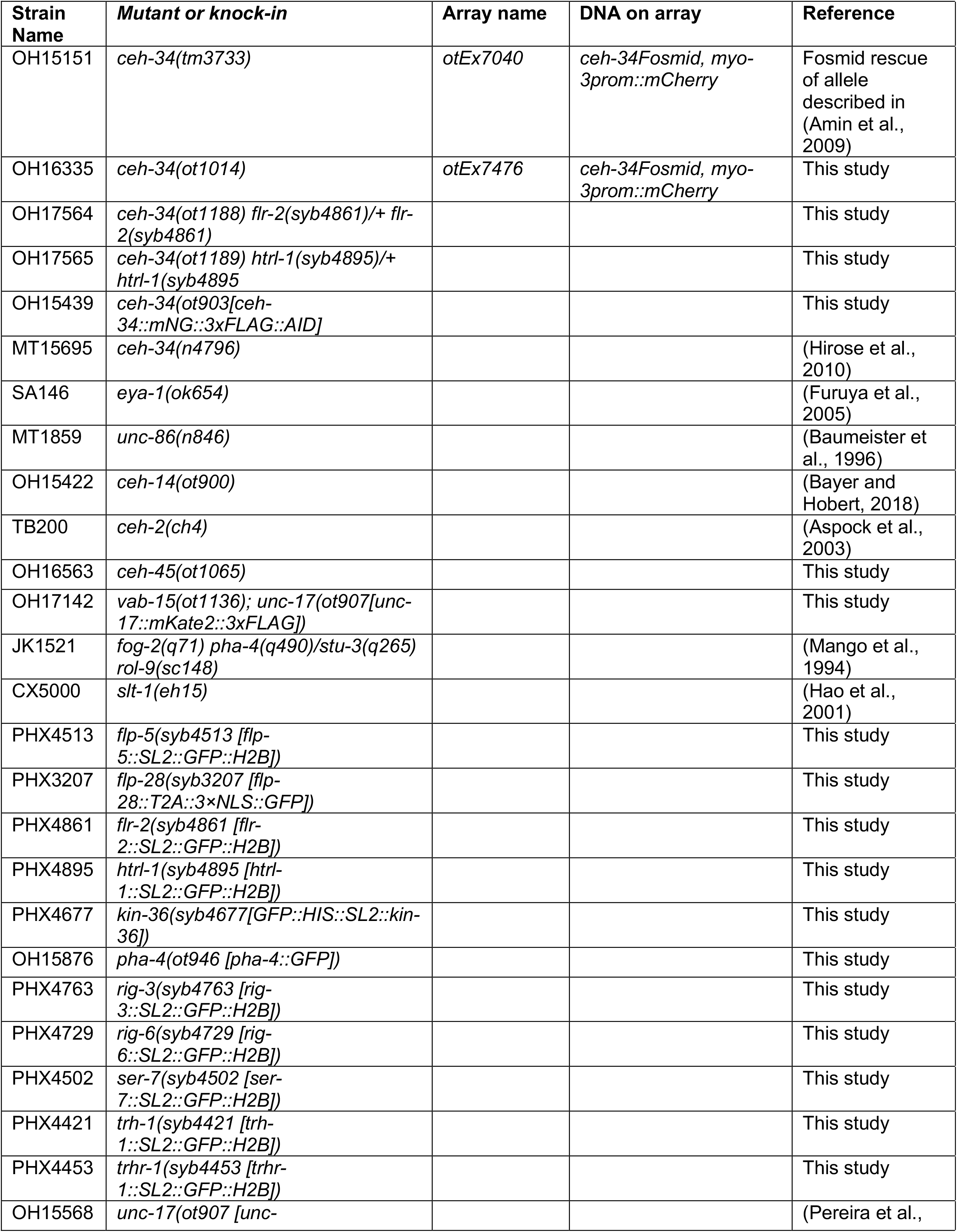
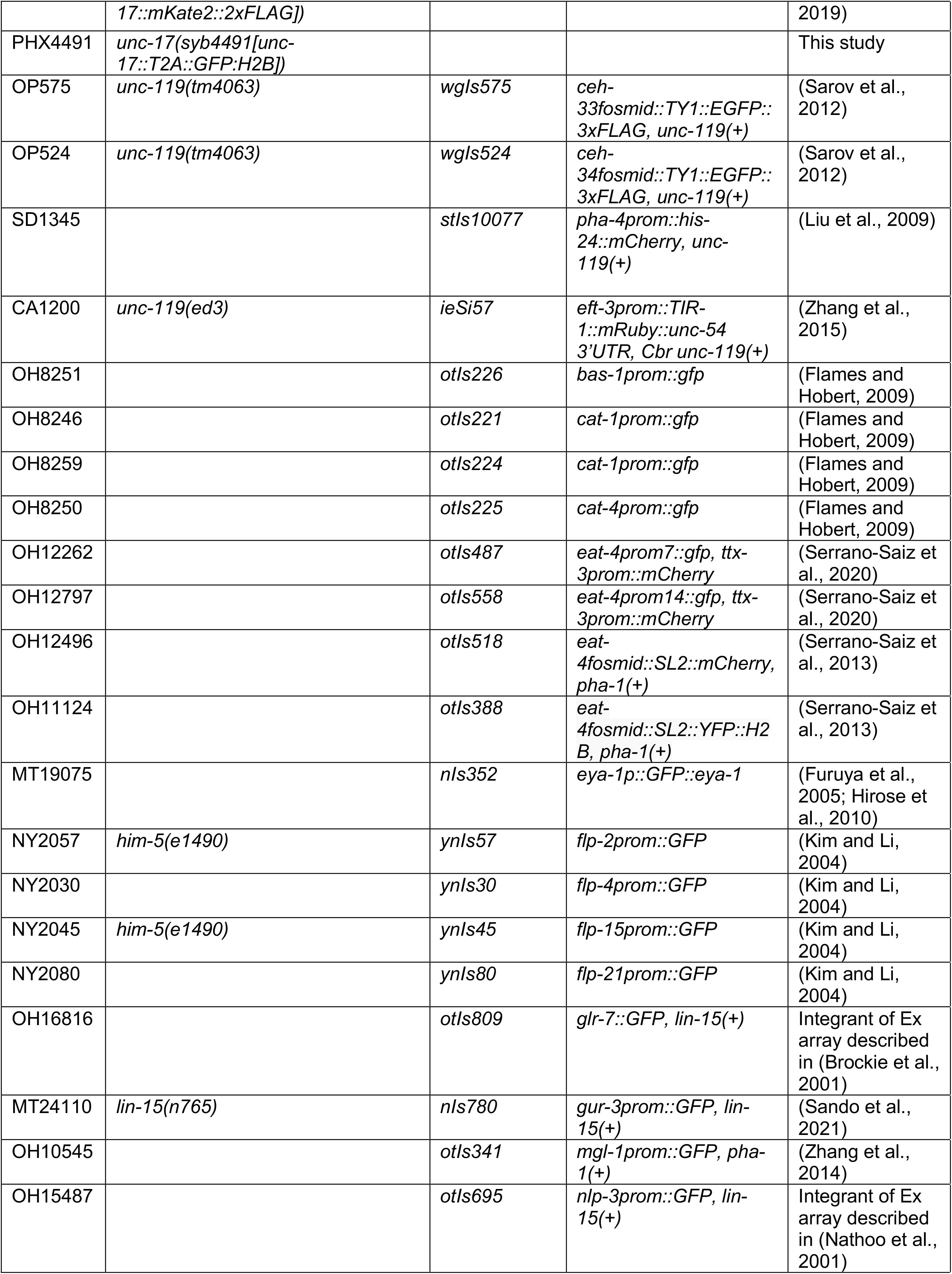
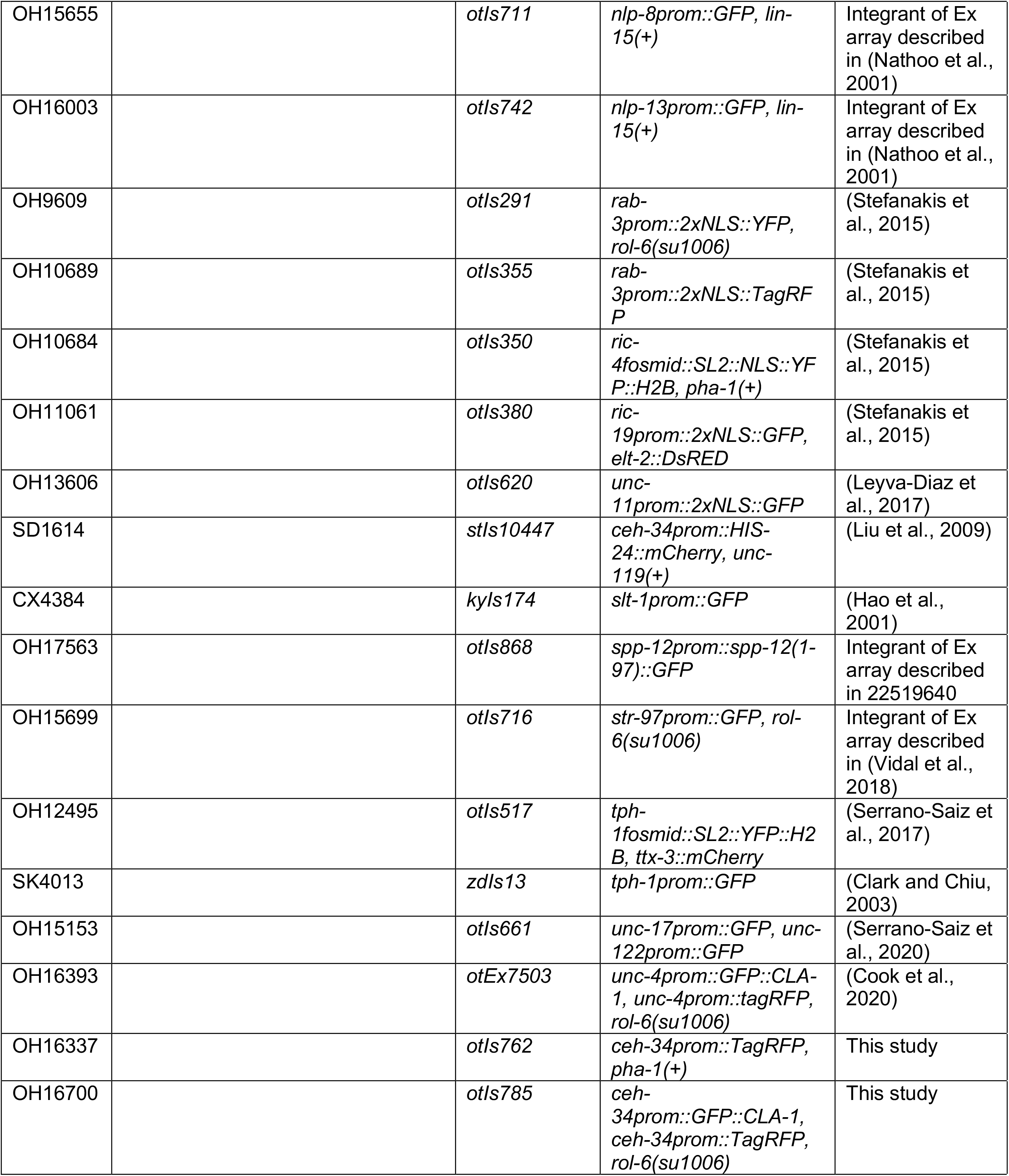

## REFERENCES

Abot, A., Cani, P.D., and Knauf, C. (2018). Impact of Intestinal Peptides on the Enteric Nervous System: Novel Approaches to Control Glucose Metabolism and Food Intake. Front Endocrinol (Lausanne) 9, 328.

Aghayeva, U., Bhattacharya, A., Sural, S., Jaeger, E., Churgin, M., Fang-Yen, C., and Hobert, O. (2021). DAF-16/FoxO and DAF-12/VDR control cellular plasticity both cell-autonomously and via interorgan signaling. BioRxiv.

Albertson, D.G., and Thomson, J.N. (1976). The pharynx of Caenorhabditis elegans. Philos Trans R Soc Lond B Biol Sci 275, 299–325.

Amin, N.M., Lim, S.E., Shi, H., Chan, T.L., and Liu, J. (2009). A conserved Six-Eya cassette acts downstream of Wnt signaling to direct non-myogenic versus myogenic fates in the C. elegans postembryonic mesoderm. Dev Biol 331, 350–360.

Arendt, D. (2008). The evolution of cell types in animals: emerging principles from molecular studies. Nature Reviews Genetics 9, 868–882.

Aspock, G., Ruvkun, G., and Burglin, T.R. (2003). The Caenorhabditis elegans ems class homeobox gene ceh-2 is required for M3 pharynx motoneuron function. Development 130, 3369–3378.

Avery, L. (2012). C. elegans feeding. WormBook, 1–23.

Avery, L., and Horvitz, H.R. (1989). Pharyngeal pumping continues after laser killing of the pharyngeal nervous system of C. elegans. Neuron 3, 473–485.

Axang, C., Rauthan, M., Hall, D.H., and Pilon, M. (2008). Developmental genetics of the C. elegans pharyngeal neurons NSML and NSMR. BMC Dev Biol 8, 38.

Ayali, A. (2004). The insect frontal ganglion and stomatogastric pattern generator networks. Neurosignals 13, 20–36.

Babu, K., Hu, Z., Chien, S.C., Garriga, G., and Kaplan, J.M. (2011). The immunoglobulin super family protein RIG-3 prevents synaptic potentiation and regulates Wnt signaling. Neuron 71, 103–116.

Banyai, L., and Patthy, L. (1998). Amoebapore homologs of Caenorhabditis elegans. Biochim Biophys Acta 1429, 259–264.

Baumeister, R., Liu, Y., and Ruvkun, G. (1996). Lineage-specific regulators couple cell lineage asymmetry to the transcription of the Caenorhabditis elegans POU gene unc-86 during neurogenesis. Genes Dev 10, 1395–1410.

Berghoff, E.G., Glenwinkel, L., Bhattacharya, A., Sun, H., Varol, E., Mohammadi, N., Antone, A., Feng, Y., Nguyen, K., Cook, S.J., et al. (2021). The Prop1-like homeobox gene unc-42 specifies the identity of synaptically connected neurons. eLife 10.

Bertani, B., and Ruiz, N. (2018). Function and Biogenesis of Lipopolysaccharides. EcoSal Plus 8.

Bhardwaj, A., Pandey, P., and Babu, K. (2020). Control of Locomotory Behavior of Caenorhabditis elegans by the Immunoglobulin Superfamily Protein RIG-3. Genetics 214, 135–145.

Bhatla, N., and Horvitz, H.R. (2015). Light and hydrogen peroxide inhibit C. elegans Feeding through gustatory receptor orthologs and pharyngeal neurons. Neuron 85, 804–818.

Brockie, P.J., Madsen, D.M., Zheng, Y., Mellem, J., and Maricq, A.V. (2001). Differential expression of glutamate receptor subunits in the nervous system of Caenorhabditis elegans and their regulation by the homeodomain protein UNC-42. J Neurosci 21, 1510–1522.

Brunet, J.F., and Pattyn, A. (2002). Phox2 genes - from patterning to connectivity. Curr Opin Genet Dev 12, 435–440.

Cheyette, B.N., Green, P.J., Martin, K., Garren, H., Hartenstein, V., and Zipursky, S.L. (1994). The Drosophila sine oculis locus encodes a homeodomain-containing protein required for the development of the entire visual system. Neuron 12, 977–996.

Colosimo, M.E., Brown, A., Mukhopadhyay, S., Gabel, C., Lanjuin, A.E., Samuel, A.D., and Sengupta, P. (2004). Identification of thermosensory and olfactory neuron-specific genes via expression profiling of single neuron types. Curr Biol 14, 2245–2251.

Cook, S.J., Crouse, C.M., Yemini, E., Hall, D.H., Emmons, S.W., and Hobert, O. (2020). The connectome of the Caenorhabditis elegans pharynx. The Journal of comparative neurology.

Cook, S.J., Jarrell, T.A., Brittin, C.A., Wang, Y., Bloniarz, A.E., Yakovlev, M.A., Nguyen, K.C.Q., Tang, L.T., Bayer, E.A., Duerr, J.S., et al. (2019). Whole-animal connectomes of both Caenorhabditis elegans sexes. Nature 571, 63–71.

Copenhaver, P.F. (2007). How to innervate a simple gut: familiar themes and unique aspects in the formation of the insect enteric nervous system. Dev Dyn 236, 1841–1864.

Corpening, J.C., Cantrell, V.A., Deal, K.K., and Southard-Smith, E.M. (2008). A Histone2BCerulean BAC transgene identifies differential expression of Phox2b in migrating enteric neural crest derivatives and enteric glia. Dev Dyn 237, 1119–1132.

Croset, V., Rytz, R., Cummins, S.F., Budd, A., Brawand, D., Kaessmann, H., Gibson, T.J., and Benton, R. (2010). Ancient protostome origin of chemosensory ionotropic glutamate receptors and the evolution of insect taste and olfaction. PLoS Genet 6, e1001064.

Dauger, S., Pattyn, A., Lofaso, F., Gaultier, C., Goridis, C., Gallego, J., and Brunet, J.F. (2003). Phox2b controls the development of peripheral chemoreceptors and afferent visceral pathways. Development 130, 6635–6642.

De Velasco, B., Shen, J., Go, S., and Hartenstein, V. (2004). Embryonic development of the Drosophila corpus cardiacum, a neuroendocrine gland with similarity to the vertebrate pituitary, is controlled by sine oculis and glass. Dev Biol 274, 280–294.

Dickinson, D.J., Pani, A.M., Heppert, J.K., Higgins, C.D., and Goldstein, B. (2015). Streamlined Genome Engineering with a Self-Excising Drug Selection Cassette. Genetics 200, 1035–1049.

Dierking, K., Yang, W., and Schulenburg, H. (2016). Antimicrobial effectors in the nematode Caenorhabditis elegans: an outgroup to the Arthropoda. Philos Trans R Soc Lond B Biol Sci 371.

Dobzhansky, T. (1964). Biology, Molecular and Organismic. American Zoologist 4, 443–452.

Dokshin, G.A., Ghanta, K.S., Piscopo, K.M., and Mello, C.C. (2018). Robust Genome Editing with Short Single-Stranded and Long, Partially Single-Stranded DNA Donors in Caenorhabditis elegans. Genetics 210, 781–787.

Dozier, C., Kagoshima, H., Niklaus, G., Cassata, G., and Burglin, T.R. (2001). The Caenorhabditis elegans Six/sine oculis class homeobox gene ceh-32 is required for head morphogenesis. Dev Biol 236, 289–303.

Drokhlyansky, E., Smillie, C.S., Van Wittenberghe, N., Ericsson, M., Griffin, G.K., Eraslan, G., Dionne, D., Cuoco, M.S., Goder-Reiser, M.N., Sharova, T., et al. (2020). The Human and Mouse Enteric Nervous System at Single-Cell Resolution. Cell 182, 1606–1622 e1623.

Du, H., and Chalfie, M. (2001). Genes Regulating Touch Cell Development in Caenorhabditis elegans. Genetics 158, 197–207.

Feng, H., and Hope, I.A. (2013). The Caenorhabditis elegans homeobox gene ceh-19 is required for MC motorneuron function. Genesis 51, 163–178.

Fung, C., and Vanden Berghe, P. (2020). Functional circuits and signal processing in the enteric nervous system. Cell Mol Life Sci 77, 4505–4522.

Furness, J.B. (2000). Types of neurons in the enteric nervous system. J Auton Nerv Syst 81, 87–96.

Furness, J.B., and Stebbing, M.J. (2018). The first brain: Species comparisons and evolutionary implications for the enteric and central nervous systems. Neurogastroenterol Motil 30.

Furuya, M., Qadota, H., Chisholm, A.D., and Sugimoto, A. (2005). The C. elegans eyes absent ortholog EYA-1 is required for tissue differentiation and plays partially redundant roles with PAX-6. Dev Biol 286, 452–463.

Ganz, J. (2018). Gut feelings: Studying enteric nervous system development, function, and disease in the zebrafish model system. Dev Dyn 247, 268–278.

Gaudet, J., and Mango, S.E. (2002). Regulation of organogenesis by the Caenorhabditis elegans FoxA protein PHA-4. Science 295, 821–825.

Gershon, M.D. (1998). The Second Brain (HarperCollins).

Gershon, M.D. (2013). 5-Hydroxytryptamine (serotonin) in the gastrointestinal tract. Curr Opin Endocrinol Diabetes Obes 20, 14–21.

Gilbert, S.F. (2019). Evolutionary transitions revisited: Holobiont evo-devo. J Exp Zool B Mol Dev Evol 332, 307–314.

Glenwinkel, L., Taylor, S.R., Langebeck-Jensen, K., Pereira, L., Reilly, M.B., Basavaraju, M., Rafi, I., Yemini, E., Pocock, R., Sestan, N., et al. (2021). In silico analysis of the transcriptional regulatory logic of neuronal identity specification throughout the C. elegans nervous system. eLife 10.

Ha, N.T., and Dougherty, K.J. (2018). Spinal Shox2 interneuron interconnectivity related to function and development. eLife 7.

Hall, D.H., and Altun, Z. (2007). C. Elegans Atlas (Cold Spring Harbor Laboratory Press).

Hanson, I.M. (2001). Mammalian homologues of the Drosophila eye specification genes. Semin Cell Dev Biol 12, 475–484.

Hao, J.C., Yu, T.W., Fujisawa, K., Culotti, J.G., Gengyo-Ando, K., Mitani, S., Moulder, G., Barstead, R., Tessier-Lavigne, M., and Bargmann, C.I. (2001). C. elegans Slit Acts in Midline, Dorsal-Ventral, and Anterior-Posterior Guidance via the SAX-3/Robo Receptor. Neuron 32, 25–38.

Hao, M.M., and Young, H.M. (2009). Development of enteric neuron diversity. J Cell Mol Med 13, 1193–1210.

Hartenstein, V. (1997). Development of the insect stomatogastric nervous system. Trends in neurosciences 20, 421–427.

Hirose, T., Galvin, B.D., and Horvitz, H.R. (2010). Six and Eya promote apoptosis through direct transcriptional activation of the proapoptotic BH3-only gene egl-1 in Caenorhabditis elegans. Proc Natl Acad Sci U S A 107, 15479–15484.

Hobert, O. (2013). The neuronal genome of Caenorhabditis elegans. WormBook, 1–106.

Hobert, O. (2016). Terminal Selectors of Neuronal Identity. Curr Top Dev Biol 116, 455–475.

Hobert, O. (2021). Homeobox genes and the specification of neuronal identity. Nat Rev Neurosci 22, 627–636.

Hobson, R.J., Hapiak, V.M., Xiao, H., Buehrer, K.L., Komuniecki, P.R., and Komuniecki, R.W. (2006). SER-7, a Caenorhabditis elegans 5-HT7-like receptor, is essential for the 5-HT stimulation of pharyngeal pumping and egg laying. Genetics 172, 159–169.

Hoeckendorf, A., Stanisak, M., and Leippe, M. (2012). The saposin-like protein SPP-12 is an antimicrobial polypeptide in the pharyngeal neurons of Caenorhabditis elegans and participates in defence against a natural bacterial pathogen. Biochem J 445, 205–212.

Horner, M.A., Quintin, S., Domeier, M.E., Kimble, J., Labouesse, M., and Mango, S.E. (1998). pha-4, an HNF-3 homolog, specifies pharyngeal organ identity in Caenorhabditis elegans. Genes Dev 12, 1947–1952.

Horvitz, H.R., Chalfie, M., Trent, C., Sulston, J.E., and Evans, P.D. (1982). Serotonin and octopamine in the nematode Caenorhabditis elegans. Science 216, 1012–1014.

Hsu, H.T., Chen, H.M., Yang, Z., Wang, J., Lee, N.K., Burger, A., Zaret, K., Liu, T., Levine, E., and Mango, S.E. (2015). TRANSCRIPTION. Recruitment of RNA polymerase II by the pioneer transcription factor PHA-4. Science 348, 1372–1376.

Hunt-Newbury, R., Viveiros, R., Johnsen, R., Mah, A., Anastas, D., Fang, L., Halfnight, E., Lee, D., Lin, J., Lorch, A., et al. (2007). High-throughput in vivo analysis of gene expression in Caenorhabditis elegans. PLoS Biol 5, e237.

Ishita, Y., Chihara, T., and Okumura, M. (2020). Serotonergic modulation of feeding behavior in Caenorhabditis elegans and other related nematodes. Neurosci Res 154, 9–19.

Jacobs, D.K., Nakanishi, N., Yuan, D., Camara, A., Nichols, S.A., and Hartenstein, V. (2007). Evolution of sensory structures in basal metazoa. Integr Comp Biol 47, 712–723.

Kalb, J.M., Lau, K.K., Goszczynski, B., Fukushige, T., Moons, D., Okkema, P.G., and McGhee, J.D. (1998). pha-4 is Ce-fkh-1, a fork head/HNF-3alpha,beta,gamma homolog that functions in organogenesis of the C. elegans pharynx. Development 125, 2171–2180.

Katidou, M., Tavernarakis, N., and Karagogeos, D. (2013). The contactin RIG-6 mediates neuronal and non-neuronal cell migration in Caenorhabditis elegans. Dev Biol 373, 184–195.

Kato, Y., Aizawa, T., Hoshino, H., Kawano, K., Nitta, K., and Zhang, H. (2002). abf-1 and abf-2, ASABF-type antimicrobial peptide genes in Caenorhabditis elegans. Biochem J 361, 221–230.

Kim, B., and Emmons, S.W. (2017). Multiple conserved cell adhesion protein interactions mediate neural wiring of a sensory circuit in C. elegans. eLife 6.

Klimovich, A., Giacomello, S., Bjorklund, A., Faure, L., Kaucka, M., Giez, C., Murillo-Rincon, A.P., Matt, A.S., Willoweit-Ohl, D., Crupi, G., et al. (2020). Prototypical pacemaker neurons interact with the resident microbiota. Proc Natl Acad Sci U S A 117, 17854–17863.

Klimovich, A.V., and Bosch, T.C.G. (2018). Rethinking the Role of the Nervous System: Lessons From the Hydra Holobiont. Bioessays 40, e1800060.

Koizumi, O. (2007). Nerve ring of the hypostome in hydra: is it an origin of the central nervous system of bilaterian animals? Brain Behav Evol 69, 151–159.

Kumar, J.P. (2009). The sine oculis homeobox (SIX) family of transcription factors as regulators of development and disease. Cell Mol Life Sci 66, 565–583.

Laranjeira, C., and Pachnis, V. (2009). Enteric nervous system development: Recent progress and future challenges. Auton Neurosci 151, 61–69.

Llewellyn-Smith, I.J. (1989). Neuropeptides and the microcircuitry of the enteric nervous system. In Regulatory Peptides, J.M. Polak, ed. (Basel: Birkhäuser Basel), pp. 247–265.

Ma, X., Zhao, Z., Xiao, L., Xu, W., Kou, Y., Zhang, Y., Wu, G., Wang, Y., and Du, Z. (2021). A 4D single-cell protein atlas of transcription factors delineates spatiotemporal patterning during embryogenesis. Nat Methods 18, 893–902.

Mackie, G.O. (1970). Neuroid conduction and the evolution of conducting tissues. Q Rev Biol 45, 319–332.

Mango, S.E. (2007). The C . elegans pharynx : a model for organogenesis. Wormbook.

Mango, S.E., Lambie, E.J., and Kimble, J. (1994). The pha-4 gene is required to generate the pharyngeal primordium of Caenorhabditis elegans. Development 120, 3019–3031.

Memic, F., Knoflach, V., Morarach, K., Sadler, R., Laranjeira, C., Hjerling-Leffler, J., Sundstrom, E., Pachnis, V., and Marklund, U. (2018). Transcription and Signaling Regulators in Developing Neuronal Subtypes of Mouse and Human Enteric Nervous System. Gastroenterology 154, 624–636.

Morarach, K., Mikhailova, A., Knoflach, V., Memic, F., Kumar, R., Li, W., Ernfors, P., and Marklund, U. (2021). Diversification of molecularly defined myenteric neuron classes revealed by single-cell RNA sequencing. Nat Neurosci 24, 34–46.

Morck, C., Rauthan, M., Wagberg, F., and Pilon, M. (2004). pha-2 encodes the C. elegans ortholog of the homeodomain protein HEX and is required for the formation of the pharyngeal isthmus. Dev Biol 272, 403–418.

Muniz, L.R., Knosp, C., and Yeretssian, G. (2012). Intestinal antimicrobial peptides during homeostasis, infection, and disease. Front Immunol 3, 310.

Myers, L., Perera, H., Alvarado, M.G., and Kidd, T. (2018). The Drosophila Ret gene functions in the stomatogastric nervous system with the Maverick TGFbeta ligand and the Gfrl co-receptor. Development 145.

Nagy, N., and Goldstein, A.M. (2017). Enteric nervous system development: A crest cell’s journey from neural tube to colon. Semin Cell Dev Biol 66, 94–106.

Narasimhan, K., Lambert, S.A., Yang, A.W., Riddell, J., Mnaimneh, S., Zheng, H., Albu, M., Najafabadi, H.S., Reece-Hoyes, J.S., Fuxman Bass, J.I., et al. (2015). Mapping and analysis of Caenorhabditis elegans transcription factor sequence specificities. eLife 4.

Ohto, H., Kamada, S., Tago, K., Tominaga, S.I., Ozaki, H., Sato, S., and Kawakami, K. (1999). Cooperation of six and eya in activation of their target genes through nuclear translocation of Eya. Mol Cell Biol 19, 6815–6824.

Ohto, H., Takizawa, T., Saito, T., Kobayashi, M., Ikeda, K., and Kawakami, K. (1998). Tissue and developmental distribution of Six family gene products. Int J Dev Biol 42, 141–148.

Oishi, A., Gengyo-Ando, K., Mitani, S., Mohri-Shiomi, A., Kimura, K.D., Ishihara, T., and Katsura, I. (2009). FLR-2, the glycoprotein hormone alpha subunit, is involved in the neural control of intestinal functions in Caenorhabditis elegans. Genes Cells 14, 1141–1154.

Patrick, A.N., Cabrera, J.H., Smith, A.L., Chen, X.S., Ford, H.L., and Zhao, R. (2013). Structure-function analyses of the human SIX1-EYA2 complex reveal insights into metastasis and BOR syndrome. Nat Struct Mol Biol 20, 447–453.

Pattyn, A., Morin, X., Cremer, H., Goridis, C., and Brunet, J.F. (1999). The homeobox gene Phox2b is essential for the development of autonomic neural crest derivatives. Nature 399, 366–370.

Pereira, L., Kratsios, P., Serrano-Saiz, E., Sheftel, H., Mayo, A.E., Hall, D.H., White, J.G., LeBoeuf, B., Garcia, L.R., Alon, U., et al. (2015). A cellular and regulatory map of the cholinergic nervous system of C. elegans. eLife 4.

Pilon, M. (2008). Fishing lines, time-delayed guideposts, and other tricks used by developing pharyngeal neurons in Caenorhabditis elegans. Dev Dyn 237, 2073–2080.

Portereiko, M.F., and Mango, S.E. (2001). Early morphogenesis of the Caenorhabditis elegans pharynx. Dev Biol 233, 482–494.

Ramakrishnan, K., and Okkema, P.G. (2014). Regulation of C. elegans neuronal differentiation by the ZEB-family factor ZAG-1 and the NK-2 homeodomain factor CEH-28. PLoS One 9, e113893.

Rao, M., and Gershon, M.D. (2018). Enteric nervous system development: what could possibly go wrong? Nat Rev Neurosci 19, 552–565.

Rauthan, M., Morck, C., and Pilon, M. (2007). The C. elegans M3 neuron guides the growth cone of its sister cell M2 via the Kruppel-like zinc finger protein MNM-2. Dev Biol 311, 185–199.

Ray, P., Schnabel, R., and Okkema, P.G. (2008). Behavioral and synaptic defects in C. elegans lacking the NK-2 homeobox gene ceh-28. Dev Neurobiol 68, 421–433.

Reilly, M.B., Cros, C., Varol, E., Yemini, E., and Hobert, O. (2020). Unique homeobox codes delineate all the neuron classes of C. elegans. Nature 584, 595–601.

Ruiz-Reig, N., Rakotobe, M., Bethus, I., Le Menn, G., Huditz, H.I., Marie, H., Lamonerie, T., and D’Autreaux, F. (2019). Developmental Requirement of Homeoprotein Otx2 for Specific Habenulo-Interpeduncular Subcircuits. J Neurosci 39, 1005–1019.

Ruvkun, G., and Hobert, O. (1998). The Taxonomy of Developmental Control in Caenorhabditis elegans. Science 282, 2033–2041.

Sasselli, V., Pachnis, V., and Burns, A.J. (2012). The enteric nervous system. Dev Biol 366, 64–73.

Schindelin, J., Arganda-Carreras, I., Frise, E., Kaynig, V., Longair, M., Pietzsch, T., Preibisch, S., Rueden, C., Saalfeld, S., Schmid, B., et al. (2012). Fiji: an open-source platform for biological-image analysis. Nat Methods 9, 676–682.

Schwarz, V., Pan, J., Voltmer-Irsch, S., and Hutter, H. (2009). IgCAMs redundantly control axon navigation in Caenorhabditis elegans. Neural Dev 4, 13.

Seo, H.C., Curtiss, J., Mlodzik, M., and Fjose, A. (1999). Six class homeobox genes in drosophila belong to three distinct families and are involved in head development. Mech Dev 83, 127–139.

Serrano-Saiz, E., Leyva-Diaz, E., De La Cruz, E., and Hobert, O. (2018). BRN3-type POU Homeobox Genes Maintain the Identity of Mature Postmitotic Neurons in Nematodes and Mice. Curr Biol 28, 2813–2823 e2812.

Serrano-Saiz, E., Poole, Richard J., Felton, T., Zhang, F., De La Cruz, Estanisla D., and Hobert, O. (2013). Modular Control of Glutamatergic Neuronal Identity in C. elegans by Distinct Homeodomain Proteins. Cell 155, 659–673.

Smit, R.B., Schnabel, R., and Gaudet, J. (2008). The HLH-6 transcription factor regulates C. elegans pharyngeal gland development and function. PLoS Genet 4, e1000222.

Sokolowski, K., Esumi, S., Hirata, T., Kamal, Y., Tran, T., Lam, A., Oboti, L., Brighthaupt, S.C., Zaghlula, M., Martinez, J., et al. (2015). Specification of select hypothalamic circuits and innate behaviors by the embryonic patterning gene dbx1. Neuron 86, 403–416.

Song, B.M., and Avery, L. (2013). The pharynx of the nematode C. elegans: A model system for the study of motor control. Worm 2, e21833.

Sulston, J.E., Schierenberg, E., White, J.G., and Thomson, J.N. (1983). The embryonic cell lineage of the nematode Caenorhabditis elegans. Dev Biol 100, 64–119.

Tadjuidje, E., and Hegde, R.S. (2013). The Eyes Absent proteins in development and disease. Cell Mol Life Sci 70, 1897–1913.

Taylor, S.R., Santpere, G., Weinreb, A., Barrett, A., Reilly, M.B., Xu, C., Varol, E., Oikonomou, P., Glenwinkel, L., McWhirter, R., et al. (2021). Molecular topography of an entire nervous system. Cell.

Tiveron, M.C., Hirsch, M.R., and Brunet, J.F. (1996). The expression pattern of the transcription factor Phox2 delineates synaptic pathways of the autonomic nervous system. J Neurosci 16, 7649–7660.

Trojanowski, N.F., Padovan-Merhar, O., Raizen, D.M., and Fang-Yen, C. (2014). Neural and genetic degeneracy underlies Caenorhabditis elegans feeding behavior. Journal of neurophysiology 112, 951–961.

Van Sinay, E., Mirabeau, O., Depuydt, G., Van Hiel, M.B., Peymen, K., Watteyne, J., Zels, S., Schoofs, L., and Beets, I. (2017). Evolutionarily conserved TRH neuropeptide pathway regulates growth in Caenorhabditis elegans. Proc Natl Acad Sci U S A 114, E4065–E4074.

Vidal, B., Aghayeva, U., Sun, H., Wang, C., Glenwinkel, L., Bayer, E.A., and Hobert, O. (2018). An atlas of Caenorhabditis elegans chemoreceptor expression. PLoS Biol 16, e2004218.

Vilimas, T., Abraham, A., and Okkema, P.G. (2004). An early pharyngeal muscle enhancer from the Caenorhabditis elegans ceh-22 gene is targeted by the Forkhead factor PHA-4. Dev Biol 266, 388–398.

Watanabe, H., Fujisawa, T., and Holstein, T.W. (2009). Cnidarians and the evolutionary origin of the nervous system. Dev Growth Differ 51, 167–183.

Wei, Z., Angerer, R.C., and Angerer, L.M. (2011). Direct development of neurons within foregut endoderm of sea urchin embryos. Proc Natl Acad Sci U S A 108, 9143–9147.

White, J.G., Southgate, E., Thomson, J.N., and Brenner, S. (1986). The structure of the nervous system of the nematode *Caenorhabditis elegans*. Philosophical Transactions of the Royal Society of London B Biological Sciences 314, 1–340.

Xuan, Z., Manning, L., Nelson, J., Richmond, J.E., Colon-Ramos, D.A., Shen, K., and Kurshan, P.T. (2017). Clarinet (CLA-1), a novel active zone protein required for synaptic vesicle clustering and release. eLife 6.

Zhang, F., Bhattacharya, A., Nelson, J.C., Abe, N., Gordon, P., Lloret-Fernandez, C., Maicas, M., Flames, N., Mann, R.S., Colon-Ramos, D.A., et al. (2014). The LIM and POU homeobox genes ttx-3 and unc-86 act as terminal selectors in distinct cholinergic and serotonergic neuron types. Development 141, 422–435.

Zhang, L., Ward, J.D., Cheng, Z., and Dernburg, A.F. (2015). The auxin-inducible degradation (AID) system enables versatile conditional protein depletion in C. elegans. Development 142, 4374–4384

## References for strain list

Bayer, E., and Hobert, O. (2018). A novel null allele of C. elegans gene ceh-14. MicroPubl Biol 2018.

Clark, S.G., and Chiu, C. (2003). C. elegans ZAG-1, a Zn-finger-homeodomain protein, regulates axonal development and neuronal differentiation. Development 130, 3781–3794.

Flames, N., and Hobert, O. (2009). Gene regulatory logic of dopamine neuron differentiation. Nature 458, 885–889.

Kim, K., and Li, C. (2004). Expression and regulation of an FMRFamide-related neuropeptide gene family in Caenorhabditis elegans. The Journal of comparative neurology 475, 540–550.

Leyva-Diaz, E., Stefanakis, N., Carrera, I., Glenwinkel, L., Wang, G., Driscoll, M., and Hobert, O. (2017). Silencing of Repetitive DNA Is Controlled by a Member of an Unusual Caenorhabditis elegans Gene Family. Genetics 207, 529–545.

Liu, X., Long, F., Peng, H., Aerni, S.J., Jiang, M., Sanchez-Blanco, A., Murray, J.I., Preston, E., Mericle, B., Batzoglou, S., et al. (2009). Analysis of cell fate from single-cell gene expression profiles in C. elegans. Cell 139, 623–633.

Nathoo, A.N., Moeller, R.A., Westlund, B.A., and Hart, A.C. (2001). Identification of neuropeptide-like protein gene families in Caenorhabditiselegans and other species. Proc Natl Acad Sci U S A 98, 14000–14005.

Pereira, L., Aeschimann, F., Wang, C., Lawson, H., Serrano-Saiz, E., Portman, D.S., Grosshans, H., and Hobert, O. (2019). Timing mechanism of sexually dimorphic nervous system differentiation. eLife 8.

Sando, S.R., Bhatla, N., Lee, E.L., and Horvitz, H.R. (2021). An hourglass circuit motif transforms a motor program via subcellularly localized muscle calcium signaling and contraction. eLife 10.

Sarov, M., Murray, J.I., Schanze, K., Pozniakovski, A., Niu, W., Angermann, K., Hasse, S., Rupprecht, M., Vinis, E., Tinney, M., et al. (2012). A genome-scale resource for in vivo tag-based protein function exploration in C. elegans. Cell 150, 855–866.

Serrano-Saiz, E., Gulez, B., Pereira, L., Gendrel, M., Kerk, S.Y., Vidal, B., Feng, W., Wang, C., Kratsios, P., Rand, J.B., et al. (2020). Modular Organization of Cis-regulatory Control Information of Neurotransmitter Pathway Genes in Caenorhabditis elegans. Genetics 215, 665–681.

Serrano-Saiz, E., Pereira, L., Gendrel, M., Aghayeva, U., Battacharya, A., Howell, K., Garcia, L.R., and Hobert, O. (2017). A Neurotransmitter Atlas of the Caenorhabditis elegans Male Nervous System Reveals Sexually Dimorphic Neurotransmitter Usage. Genetics 206, 1251–1269.

Stefanakis, N., Carrera, I., and Hobert, O. (2015). Regulatory Logic of Pan-Neuronal Gene Expression in C. elegans. Neuron 87, 733–750.

